# CD19-CAR T cells reset the EBV^+^ autoreactive B cell repertoire in autoimmune disease

**DOI:** 10.64898/2026.10.24.761515

**Authors:** Ya’el Courtney, Mahesh Pandit, Laura S. van Dam, Shady Younis, Sajede Rasouli, Mengrui Zhang, Yashaar Chaichian, Shima Parsafar, Tobias V. Lanz, Eline J. Arends, Cees van Kooten, Y.K. Onno Teng, Melanie Hagen, Andreas Wirsching, Fabian Müller, Andreas Mackensen, Aline Bozec, Ricardo Grieshaber-Bouyer, Georg Schett, William H. Robinson

**Affiliations:** Division of Immunology and Rheumatology, Department of Medicine, Stanford University School of Medicine, Stanford, CA, USA; VA Palo Alto Health Care System, Palo Alto, CA, USA; Quantitative Sciences Unit, Stanford University School of Medicine, Stanford, CA, USA; Institute for Immunity, Transplantation and Infection, Stanford University School of Medicine, Stanford, CA, USA; Lupus, Vasculitis and Complement-associated Systemic Diseases (LuVaCs), Department of Nephrology, Leiden University Medical Center, Leiden, the Netherlands; Department of Internal Medicine 3 – Rheumatology and Immunology, Friedrich-Alexander-Universität Erlangen-Nürnberg and Universitätsklinikum Erlangen, Erlangen, Germany; Department of Internal Medicine 5 – Hematology and Oncology, Friedrich-Alexander-Universität Erlangen-Nürnberg and Universitätsklinikum Erlangen, Erlangen, Germany; Deutsches Zentrum Immuntherapie (DZI), Friedrich-Alexander-Universität Erlangen-Nürnberg and Universitätsklinikum Erlangen, Erlangen, Germany

## Abstract

Epstein-Barr virus (EBV) reprograms autoreactive B cells into activated antigen-presenting cells that drive pathogenic T- and B-cell responses in autoimmune disease.^1–4^ Whether eliminating these cells contributes to durable remission after CD19 chimeric antigen receptor (CAR) T-cell therapy^5–7^ is unknown. Here, using EBV B-cell sequencing (EBV-seq)^3^, we profiled EBV^+^ B cells from patients treated with CD19-CAR T cells (systemic lupus erythematosus, n=6; inflammatory myopathy, n=3; systemic sclerosis, n=3) or rituximab (n=15) and recombinantly expressed their antibodies to define antigen specificity. At baseline, EBV^+^ B cells were enriched relative to healthy donors, class-switched and hypermutated, expressed the latency III program with heightened antigen-presentation signatures, and encoded autoantibodies. CAR T cells, but not rituximab, depleted EBV^+^ B cells and shifted the reconstituted repertoire towards IgM^+^, minimally mutated, largely non-autoreactive cells, consistent with infection of newly generated B cells. After rituximab, EBV^+^ B cells retained a class-switched, hypermutated and autoreactive with an activated antigen-presenting phenotype. Sustained drug-free remission occurred in 8/12 CAR T-cell-treated versus 0/15 rituximab-treated patients. *In vitro*, EBV^+^ B cells resisted anti-CD20 and anti-CD19 antibodies but were killed by CAR T cells and a CD19 T-cell engager. Our findings suggest resetting the EBV^+^ autoreactive B-cell repertoire contributes to sustained drug-free remission.

---

Chimeric antigen receptor (CAR) T cells have emerged as a transformative therapeutic modality for autoimmune diseases.^5,7,8^ In contrast to B cell depletion with anti-CD20 or anti-CD19 antibodies, which provide benefit but rarely result in long-term drug-free remission^9,10^, CD19-CAR T-cell therapy induces sustained drug-free remission in patients with systemic lupus erythematosus (SLE), idiopathic inflammatory myopathies (IIM) and systemic sclerosis (SSc).^8,11^ A systematic review of 145 patients across 16 studies reported that 84% of CD19-CAR T-cell-treated SLE patients were in drug-free remission at 1 year.^12^ The approach has since been extended to additional autoimmune diseases^7,11^ as well as to other modalities including *in vivo* CAR T-cell generation by mRNA-lipid nanoparticle or lentiviral delivery^13,14^ and T-cell engager bi-specific antibodies^15–18^. The durability of these responses has made the mechanism of immune reset a central question for the field.^19,20^

CD19-CAR T-cell therapy clears CD19⁺ and CD20⁺ B cells from secondary lymphoid organs and non-lymphoid tissue, and additionally disrupts follicular architecture with loss of follicular dendritic cells, while sparing plasma cells, T cells and macrophages.^21^ Antibody-based depletion behaves differently: rituximab and obinutuzumab reduce but do not reliably eliminate lymph-node B cells and leave follicular structures intact, despite achieving comparable depletion of circulating B cells.^21^ Following CAR T-cell therapy, B cells reconstitute at approximately 3-4 months post-CD19-CAR T from naïve, non-class-switched precursors, typically without disease relapse.^5,8^ Whether sustained remission requires elimination of a specific pathogenic B-cell population, a more complete depletion of all B cells, or disruption of the follicular niche that sustains them, remains unresolved.

Epstein-Barr virus (EBV) infection is strongly associated with SLE, multiple sclerosis (MS) and other autoimmune diseases.^1,2,22,23^ EBV establishes lifelong latency in the memory B cell compartment^24^, which places the virus in precisely the cell type that B cell depleting therapies target. Several mechanisms have been proposed that link that latency to autoimmunity, and they are not mutually exclusive. Molecular mimicry between EBV nuclear antigen 1 (EBNA1) and central nervous system autoantigens, including GlialCAM and anoctamin-2, generates cross-reactive antibody and T cell responses in MS.^25,26^ Functional mimicry by latent membrane protein 1 (LMP1), a constitutively active CD40 mimic, delivers survival and activation signals to B cells that would otherwise be deleted.^27^ EBNA2 binds at autoimmune risk loci across SLE and multiple other autoimmune diseases, placing a viral transcription factor directly on the genetic architecture of autoimmunity.^28^ Lytic reactivation expands the infected pool and the antigen load^29^, and impaired cytotoxic control of EBV is documented in both SLE and MS.^30,31^ Early autoantibody responses in lupus initiate against EBNA1 epitopes before spreading to self^32^, and antibodies to EBV antigens cross-react with lupus autoantigens.^1,22^

Recently, we demonstrated that EBV directly infects autoreactive B cells in SLE and MS and, through expression of the EBV transcription factor EBNA2, reprograms these cells into activated antigen-presenting B cells capable of driving systemic autoreactive T and EBV^-^ B cell responses that underlie the pathogenesis of autoimmune diseases.^3,4^ This model predicts that the EBV^+^ autoreactive B cell is the population whose elimination determines whether an immune reset holds. On the other hand, therapies which spare these cells will not produce durable remission. This concept has not been tested in patients.

Here, we investigated the impact of CD19-CAR T-cell therapy, as compared with the anti-CD20 B cell depleting antibody rituximab, on EBV^+^ B cells in patients with SLE, IIM and SSc. We utilized EBV-single cell RNA-sequencing (EBV-seq) to identify and characterize EBV^+^ B cells at baseline and following treatment, and to define the transcriptional programs and EBV latency states of EBV^+^ B cells, and the antibody specificities they encode. In addition, we characterized the susceptibility of EBV^+^ B cells to killing by monoclonal antibodies and T cell-mediated modalities, including CD19-CAR T-cells and T-cell engagers.

## Results

### Patient characteristics

Using a refined version of EBV-seq^3^ (EBV-seq.2) we profiled 295,382 B cells from 12 patients with autoimmune disease treated with CD19-CAR T-cells (SLE, n=6; IIM, n=3; SSc, n=3)^7^ and 15 patients with SLE treated with B-cell-depleting antibody therapy (Table 1, Fig. 1a). The latter comprised 7 patients who received rituximab plus belimumab in the Synbiose-II trial and contributed paired samples at week 0 and week 44 (Supplementary Table 3), and 8 patients who received rituximab and contributed a single sample collected at least 6 months after infusion (Supplementary Table 2).

**Table 1.** Characteristics of 27 Patients with Autoimmune Disease Treated with CD19-CAR T Cell Therapy or B Cell-Depleting Antibody Therapy.

|  | **CD19-CAR T<br>SLE (N=6)** | **CD19-CAR T<br>IIM (N=3)** | **CD19-CAR T<br>SSc (N=3)** | **Rituximab<br>SLE (N=8)** | **Rituximab +<br>belimumab<br>SLE (N=7)** |
|---|---|---|---|---|---|
| **Age, years** | 33 (21-56) | 61 (45-81) | 36 (23-56) | 37 (23-62) | 35 (17-46) |
| **Sex, females, N (%)** | 3/6 (50%) | 2/3 (67%) | 1/3 (33%) | 8/8 (100%) | 6/7 (86%) |
| **Disease duration, years** | 6.5 (1-25) | 1.3 (1-2) | 3 (1-4) | 10* (0-21) | 8.6 (0-26) |
| **Disease activity, units** | SLEDAI: 8 (6;11) | MMT8: 133 (74;143)<br>FVC: 3.4 (3.1;3.8) L | mRSS: 26 (24;35)<br>FVC: 2.8 (1.9;2.8) L | SLEDAI-2K: 8 (0.25;12) | SLEDAI-2K: 22 (21;23.5) |
| **Prior therapies, N** | 5 (3.75;6.75) | 2 (2;2) | 3 (3;4) | 5.5 (5;7) | 5 (3;5) |
| **Clinical response, N** | DORIS remission (6/6) | Moderate/major ACR response (3/3) | CRISS-25 (3/3) | DORIS remission (2/8) | Clinical remission on therapy (7/7); LLDAS (6/7) |
| **Initial drug-free state, N** | 6/6 | 3/3 | 3/3 | 0/8 | 0/7 |
| **Sustained drug-free state, N** | 5/6 | 2/3 | 1/3 | 0/8 | 0/7 |
| **Duration of DF state<sup>+</sup>, months** | 31.7 (23.2;40.8) | 24.0 (10.3;31.3) | 24.4 (22.9;29.9) | N.A. | N.A. |
| **Relapse or treatment failure, N** | 1/6 | 1/3 | 2/3 | 6/8 | 4/7‡ |
| **Time to relapse, months** | 16.6 (9.3;28.2) |  |  | N.A. | 8.3 (6.4;13.6)‡ |
| **Sampling timepoints** | -1 month<br>+6 months | -1 month<br>+6 months | -1 month<br>+6 months | +6 months | Week 0<br>Week 44 |

**Values are medians (interquartile ranges) unless otherwise specified. Age is presented as mean (range). Disease activity is given at baseline for the CD19-CAR T and rituximab + belimumab groups and at the time of post-treatment sampling for the rituximab group. SLE, systemic lupus erythematosus; IIM, idiopathic inflammatory myopathies; SSc, systemic sclerosis; FVC, forced vital capacity; SLEDAI-2K, Systemic Lupus Erythematosus Disease Activity Index-2000; MMT8, Manual Muscle Testing 8; mRSS, modified Rodnan skin score; CRISS, Composite Response Index in Systemic Sclerosis; DORIS, Definitions Of Remission In SLE; LLDAS, Lupus Low Disease Activity State; DF, drug-free; N.A., not applicable.**
**DORIS remission is an on-therapy definition. All 12 CD19-CAR T recipients were off immunosuppressive therapy at the time of response assessment, whereas every patient treated with B cell-depleting antibody therapy remained on prednisone and/or maintenance immunosuppression; no patient in either rituximab-treated group achieved drug-free remission. DORIS is defined for SLE only and is therefore not applicable to the IIM and SSc groups.**
**\*Disease duration in years for the rituximab SLE patients is calculated using 7 of 8 patients, as the exact duration is unclear for patient 1.**
**<sup>+</sup>time from CAR T-cell infusion until last follow-up or relapse.**
**‡Protocol-defined treatment failure in the Synbiose-II trial, at weeks 28, 28, 44 and 104; converted to months for comparability.**

**Fig. 1.**
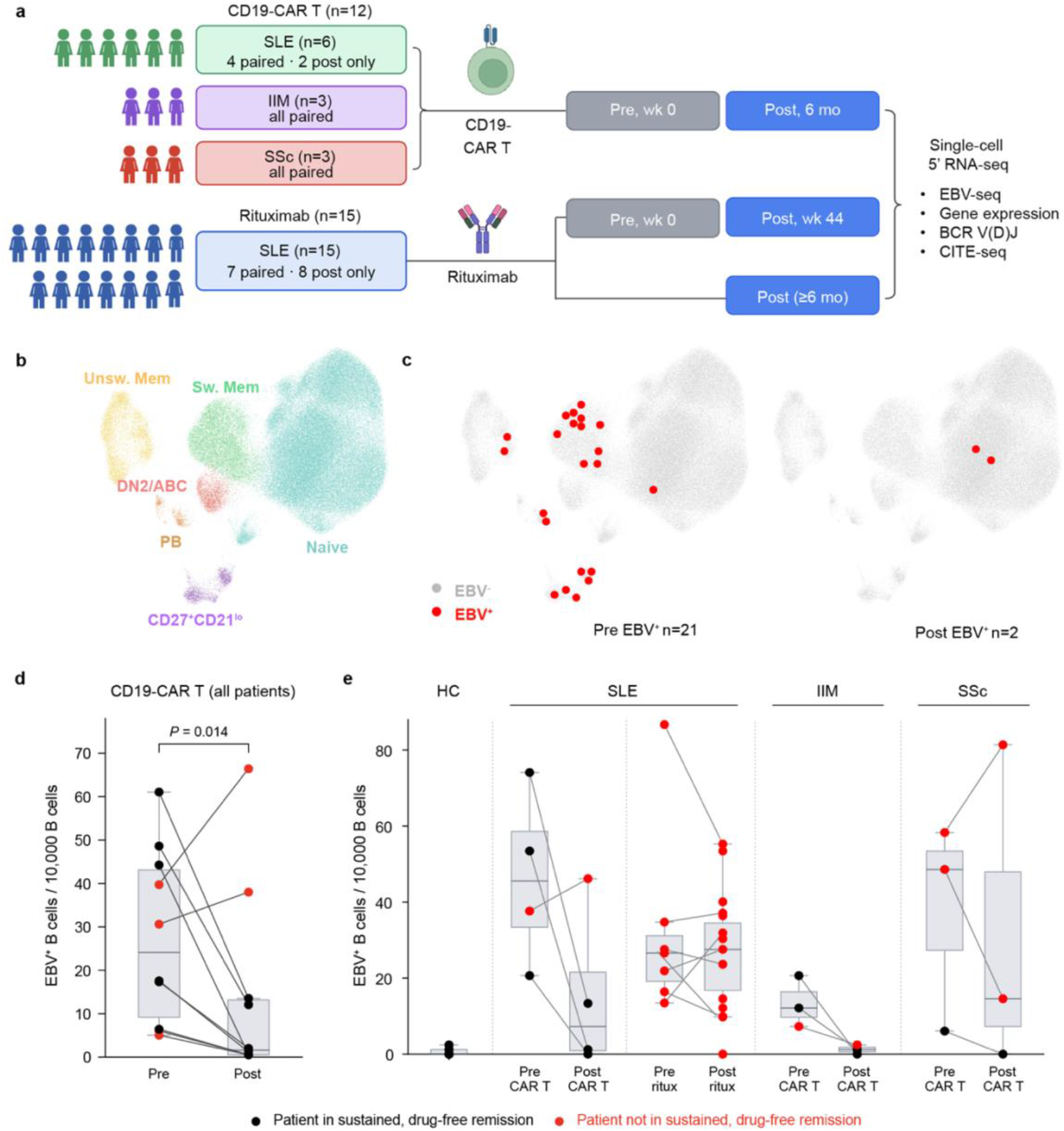
Frequency of EBV⁺ B cells before and after CD19-CAR T-cell therapy or rituximab. **a**, Study design. Peripheral-blood B cells were profiled by EBV-seq (EBV-seq.2) in 12 patients with autoimmune disease treated with CD19-CAR T cells (SLE, n = 6, of whom 4 contributed paired pre- and post-treatment samples and 2 contributed post-treatment samples only; IIM, n = 3, all paired; SSc, n = 3, all paired) and in 15 patients with SLE treated with rituximab-based B cell depletion (7 with paired week 0 and week 44 samples, 8 with a post-treatment sample only collected ≥6 months after infusion). Gene expression, BCR V(D)J, surface protein and EBV-seq data were obtained from the same cells in every library. **b**, UMAP embedding of 295,382 B cells, colored by B-cell subset. **c**, EBV⁺ B cells (red) projected onto the UMAP embedding before and after CD19-CAR T-cell therapy in one representative patient with SLE; gray, EBV⁻ B cells. N indicates the number of EBV⁺ B cells detected at each time point. **d**, Frequency of EBV⁺ B cells (per 10,000 B cells) before and after CD19-CAR T-cell therapy, all patients pooled; analysis restricted to the 10 patients with paired pre- and post-treatment samples (SLE, n = 4; IIM, n = 3; SSc, n = 3); two-sided Wilcoxon signed-rank test. **e**, Frequency of EBV⁺ B cells (per 10,000 B cells) in healthy controls (HC) and by disease group and treatment. Exact P values above each group are for the comparison with healthy controls, in bold where P < 0.05 and in gray otherwise. In **d** and **e**, black denotes patients in sustained drug-free remission and red denotes patients not in sustained drug-free remission; horizontal lines within the boxes indicate medians and the boxes indicate interquartile ranges. Whiskers extend to the most extreme value within 1.5 times the interquartile range of the nearest quartile. Comparisons with healthy controls in e are two-sided Mann-Whitney U tests.

CD19-CAR T cells achieved initial drug-free remission in 6/6 patients with SLE, moderate or major responses in 3/3 patients with IIM, and clinical responses in 3/3 patients with SSc; 8/12 patients were in sustained drug-free remission at a median follow-up for 31 months (Table 1; Supplementary Table 1). One patient with SLE, one with IIM and two with SSc relapsed at months 9, 10, 23 and 30 after CD19-CAR T-cell infusion, respectively. In contrast, 0/15 patients treated with the B-cell-depleting antibody therapy rituximab achieved drug-free remission (Supplementary Table 2).

### EBV-seq analysis of the B cell repertoire

Uniform Manifold Approximation and Projection (UMAP) analysis of the integrated B cell transcriptome revealed distinct subtype populations spanning the naïve-to-effector differentiation spectrum (Fig. 1b; Extended Data Fig. 1a-b, 2a). EBV^+^ B cells were detected in all patients at baseline across SLE, IIM and SSc, and localized predominantly to activated and memory B cell compartments in pre-treatment samples (Fig. 1c; Extended Data Fig. 2b).

EBV⁺ B cell frequency was significantly reduced 6 months after CD19-CAR T-cell therapy across diseases (Fig. 1d; n = 10 patients with paired samples; pooled two-sided Wilcoxon signed-rank P=0.014), whereas rituximab-treated patients retained EBV⁺ B cells at frequencies comparable to pre-treatment levels (Fig. 1e). Healthy controls carried significantly fewer EBV⁺ B cells than patients in every autoimmune disease group before treatment (median 0.4 per 10,000 B cells; Fig. 1e).

In the patient with SSc (#9) who relapsed, EBV^+^ B cell frequency was relatively high at baseline and did not decrease after treatment, and two EBV^+^ clonotypes persisted across time points (Extended Data Fig. 3a). No shared EBV^+^ clonotypes were detected before and after CD19-CAR T-cell therapy in any other patient.

### Analysis of B cell phenotype

EBV^+^ B cells prior to CD19-CAR T-cell therapy were predominantly class-switched (IgG and IgA) in SLE and IIM, with elevated somatic hypermutation (SHM) rates, consistent with antigen-experienced, germinal center–derived populations (Fig. 2a-c). Following CD19-CAR T-cell therapy, EBV⁺ B cells shifted toward IgM-dominant isotype usage in SLE (31% to 0% class-switched) and IIM (53% to 28%), accompanied by reduced SHM in SLE (P<0.0001), IIM (P<0.0001) and SSc (*P*=0.0027) (Fig. 2c). This is consistent with new infection of naïve B cells by EBV rather than regeneration of an EBV^+^ B cell pool from infected memory B cells surviving CD19-CAR T-cell therapy.

**Fig. 2.**
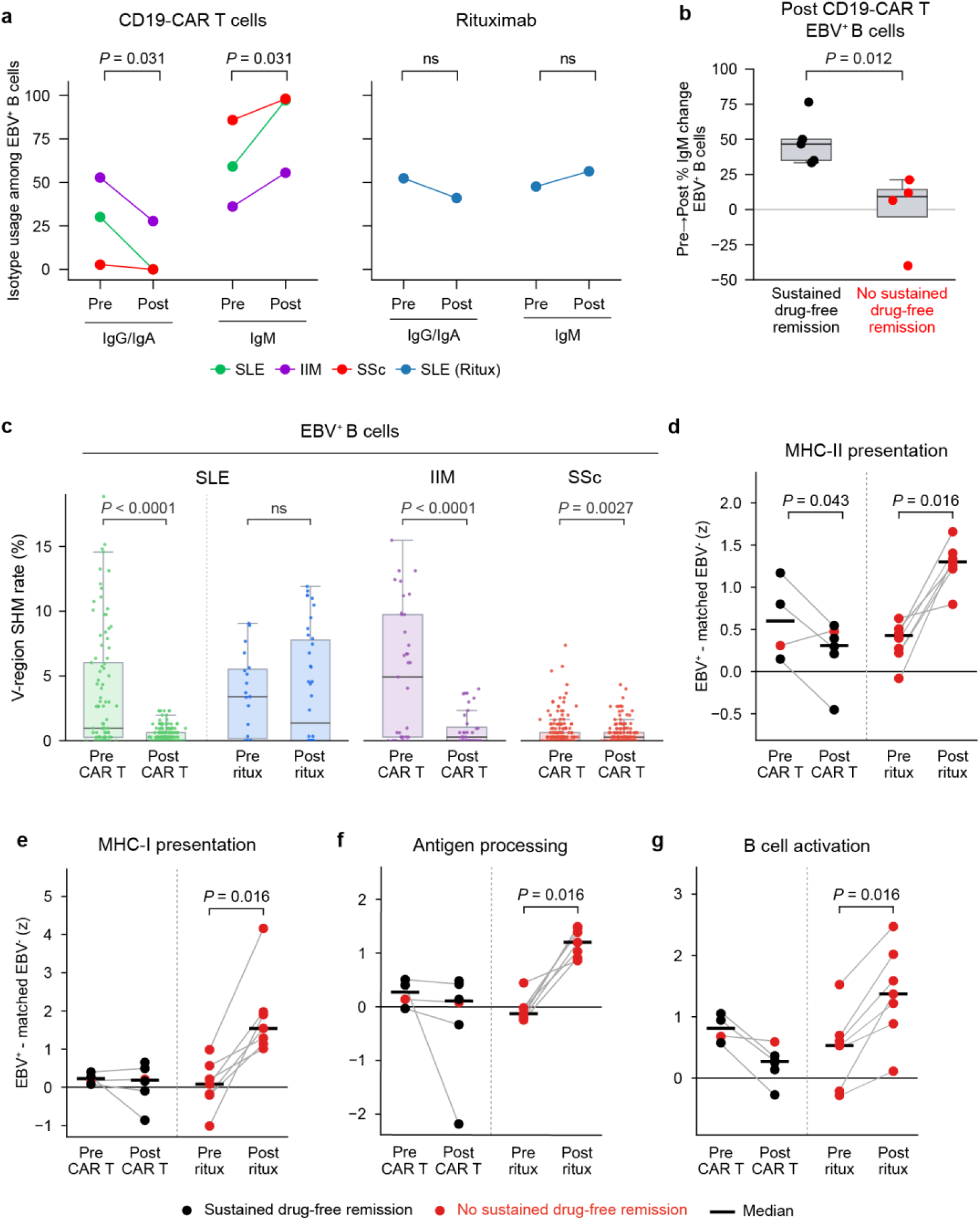
CD19-CAR T-cell therapy resets the EBV⁺ B-cell repertoire toward a naïve phenotype. **a**, Percentage of class-switched (IgG⁺ or IgA⁺, left) and IgM⁺ (right) EBV⁺ B cells before and after CD19-CAR T-cell therapy in SLE, IIM and SSc, and before and after rituximab in SLE. Each point is the pooled value for one disease group at one time point and lines connect time points within a group; P values for the change in class-switched fraction after CD19-CAR T-cell therapy are shown for SLE and IIM, and the SSc and rituximab comparisons are marked ns. **b**, Pre- to post-treatment change in the percentage of IgM⁺ cells among EBV⁺ B cells (left) and among EBV⁻ B cells (right) after CD19-CAR T-cell therapy, in patients who achieved sustained drug-free remission (black) and patients who did not (red); analysis restricted to the 10 patients with paired samples. Numbers identify individual patients; two-sided Mann–Whitney U test. **c**, V-region somatic hypermutation rate of EBV⁺ B cells by disease group and treatment. Each dot represents one cell; bars show the median and interquartile range. P values are cell level. **d–g**, Module scores for MHC class II (d), MHC class I (e), antigen processing (f), and B cell activation (g) in EBV⁺ B cells, expressed as the difference from depth-matched EBV⁻ B cells from the same library (z-score). Each point is one patient, lines connect paired samples and horizontal bars indicate medians; two-sided Wilcoxon signed-rank tests.

This reset of the EBV^+^ B cells was associated with clinical outcome. Patients who achieved sustained drug-free remission showed a greater pre- to post-treatment increase in the IgM fraction of EBV^+^ B cells than patients who did not (Fig. 2b; n = 10 patients with paired samples; *P*=0.02), with no corresponding difference in the EBV^-^ compartment (*P*=0.58). A comparable shift toward unmutated IgM was seen among EBV^-^ B cells overall (Extended Data Fig. 4a-b), consistent with reconstitution of the whole B cell pool from naïve precursors; the association with sustained remission, however, was confined to the EBV^+^ compartment.

In contrast, EBV⁺ B cells from rituximab-treated patients with SLE retained class-switched isotypes, and SHM did not differ between pre- and post-rituximab samples (*P*=0.88; Fig. 2c), indicating failure to reset the EBV⁺ B cell repertoire. Relative to depth-matched EBV⁻ B cells from the same libraries, EBV⁺ B cells after rituximab showed increased MHC class II, MHC class I, antigen processing, and B cell activation module scores (Fig. 2d-g; all 7 patients increased in every module, Wilcoxon signed-rank *P* = 0.016), with no comparable change across CD19-CAR T-cell therapy.

### Autoreactivity of antibodies encoded by EBV^+^ B cells

To determine whether EBV^+^ B cells encode autoreactive antibodies, we recombinantly expressed 50 monoclonal antibodies (mAbs) from individual EBV^+^ B cells and assessed reactivity by HEp-2 immunofluorescence and ELISA (Fig. 3). Pre-treatment EBV^+^ B cells from patients with SLE encoded mAbs with diverse autoreactive staining patterns, including nuclear homogeneous, cytoplasmic, nucleolar, and mitotic patterns, and 12 of 14 (86%) were HEp-2 reactive (Fig. 3a, c; Supplementary Fig. 1).

**Fig. 3.**
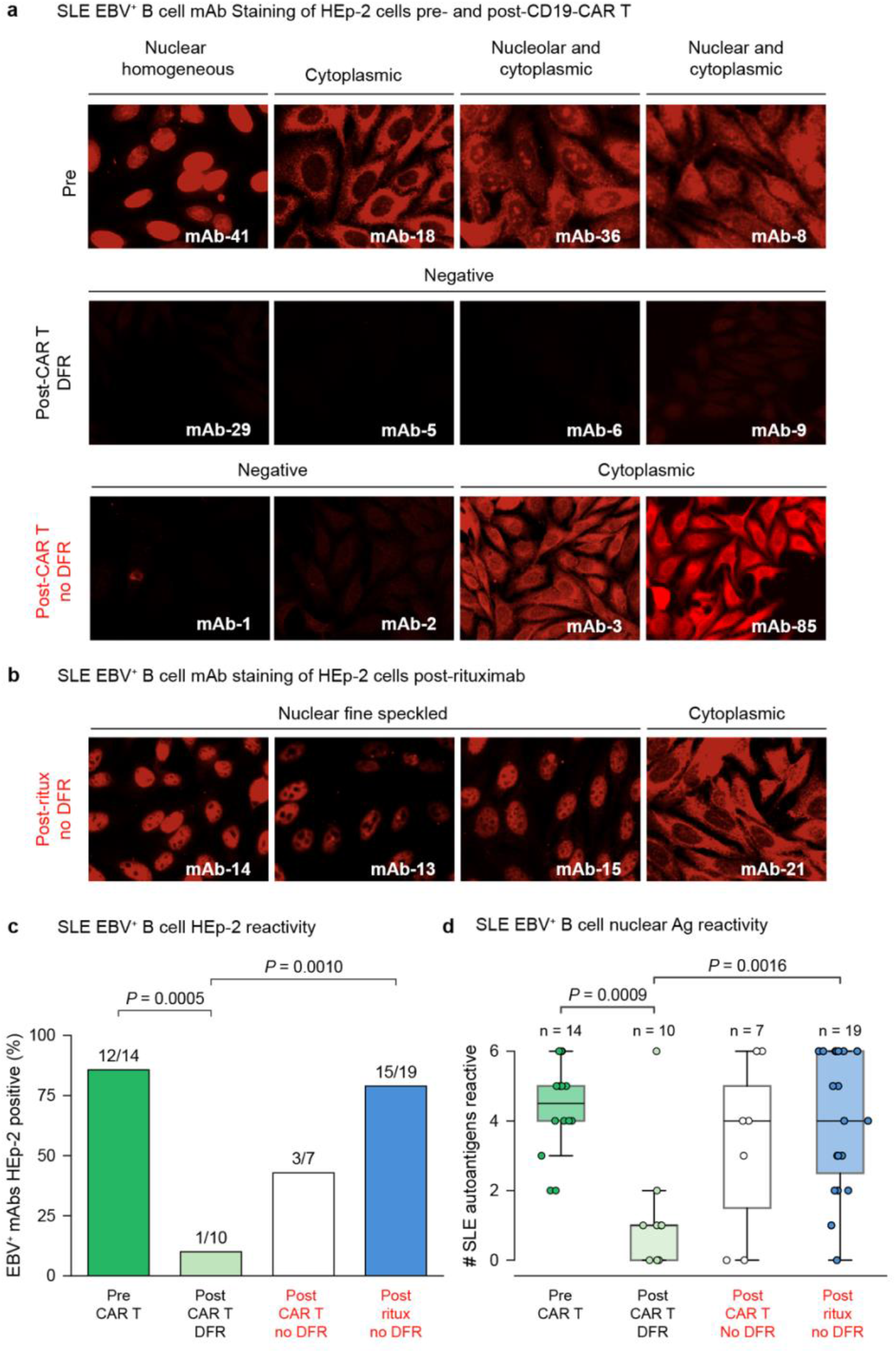
Autoreactivity of monoclonal antibodies derived from EBV⁺ B cells. **a**, Representative HEp-2 indirect immunofluorescence staining of monoclonal antibodies (mAbs) recombinantly expressed from single EBV⁺ B cells of patients with SLE before CD19-CAR T-cell therapy (top), after CD19-CAR T-cell therapy in patients in sustained drug-free remission (DFR, middle), and after CD19-CAR T-cell therapy in the patient not in sustained DFR (bottom). Staining patterns are indicated above each group of images. **b**, HEp-2 staining of mAbs cloned from EBV⁺ B cells after rituximab treatment; none of these patients were in DFR. **c**, Percentage of HEp-2-positive mAbs by treatment group: 12 of 14 (86%) before CD19-CAR T-cell therapy, 1 of 10 (10%) after CD19-CAR T-cell therapy in sustained DFR, 3 of 7 (43%) after CD19-CAR T-cell therapy without sustained DFR, and 15 of 19 (79%) after rituximab; two-sided Fisher’s exact test. **d**, Number of SLE-associated nuclear autoantigens recognized per mAb by ELISA in the same four groups; n indicates the number of mAbs per group. Each dot represents one mAb, horizontal lines within the boxes indicate medians and the boxes indicate interquartile ranges; two-sided Mann–Whitney U test. The remaining EBV⁺ mAbs are shown in Supplementary Fig. 1 and the EBV⁻ mAbs of the same patients in Extended Data Fig. 3 and Supplementary Fig. 2.

After CD19-CAR T-cell therapy, HEp-2 reactivity fell to 1 of 10 mAbs (10%) in patients in sustained drug-free remission (*P*=0.0005; Fig. 3c). In the patient with SLE who did not achieve sustained drug-free remission, 3 of 7 post-treatment mAbs (43%) remained reactive, among them mAb-3, an unmutated IgM antibody with cytoplasmic and nuclear staining cloned from an EBV^+^ B cell in latency III (Extended Data Fig. 3b).

Post-rituximab EBV^+^ B-cell-derived mAbs were HEp-2 reactive more often (15 of 19, 79%) than those from CAR T-cell-treated patients in sustained drug-free remission (1 of 10, 10%; *P*=0.0010) (Fig. 3b, c; Supplementary Fig. 1). The same ordering held for the breadth of nuclear autoantigen recognition by ELISA (Fig. 3d; *P*=0.0009 for pre-treatment vs. post-CD19-CAR T-cell therapy in remission, *P*=0.0016 for post-rituximab versus post-CD19-CAR T-cell therapy in remission).

These frequencies can be compared to those in the EBV⁻ repertoire of the same patients, in which 6 of 40 pre-treatment mAbs (15%) and 1 of 10 post-treatment mAbs (10%) were HEp-2 reactive (Extended Data Fig. 3c-g; Supplementary Fig. 2). Autoreactivity was therefore enriched in the EBV⁺ compartment before treatment, and after CD19-CAR T cell therapy the EBV⁺ compartment returned to the background frequency of the EBV⁻ compartment.

### EBV^+^ B cells are predominantly in latency III and carry its transcriptional signature

EBV⁺ B cells from healthy EBV-seropositive donors were entirely in latency 0/I, consistent with the resting memory reservoir that sustains lifelong carriage. In contrast, a large fraction of EBV⁺ B cells across SLE, IIM, and to a lesser degree SSc expressed viral genes characteristic of latency III (Fig. 4a and Extended Data Fig. 5a). The distribution of latency programs remained stable from before to after CD19-CAR T-cell therapy, even though the post-treatment EBV⁺ B cells arose from newly infected naïve cells. EBV⁺ B cells surviving rituximab were predominantly in latency III. Latency III is likely a property of the infected state in autoimmune disease rather than of the chronically infected memory pool, and it persists through both forms of B cell depletion.

**Fig. 4.**
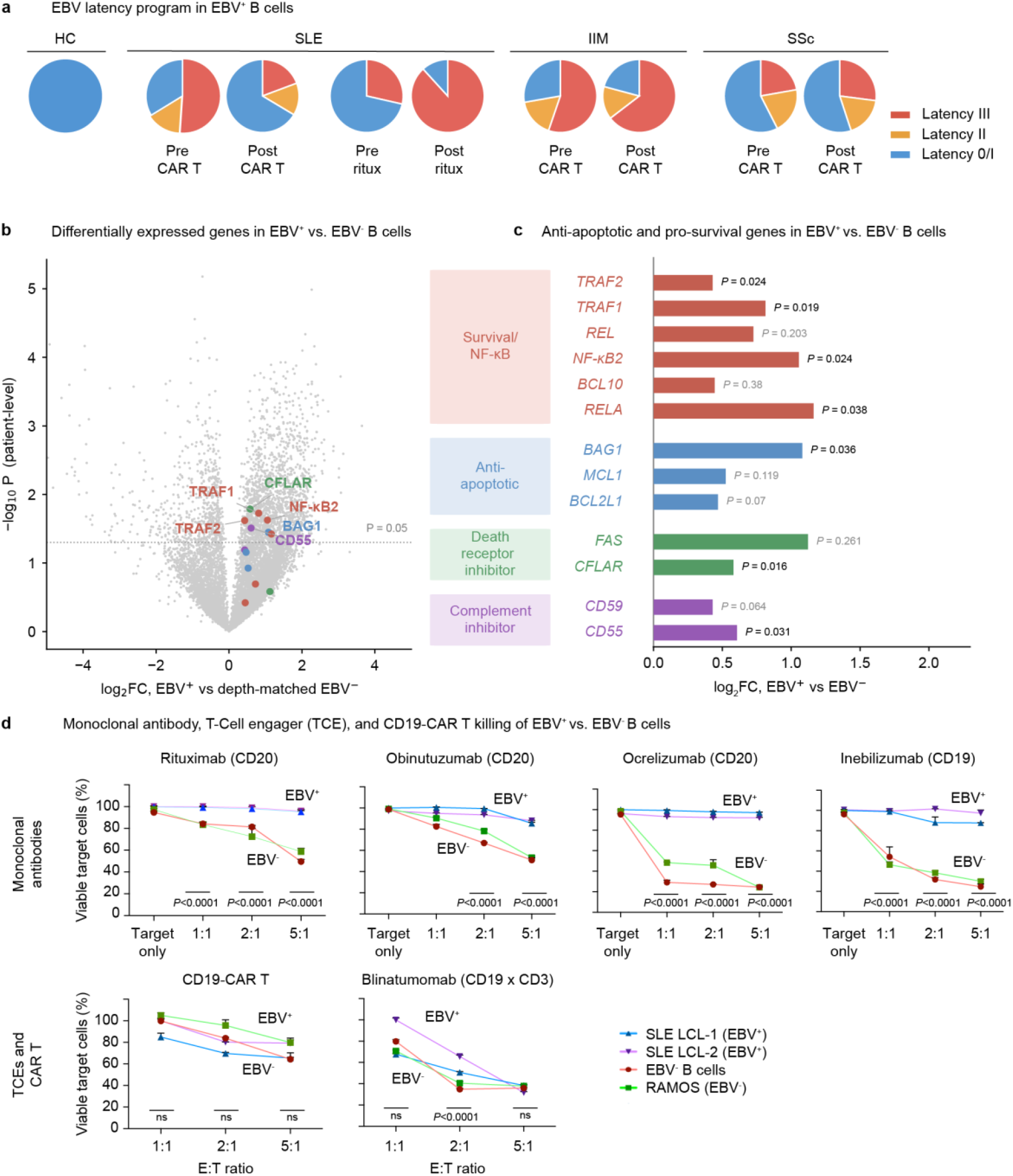
EBV latency programs and susceptibility of EBV⁺ B cells to antibody- and T-cell-mediated cytotoxicity. **a**, Distribution of EBV latency programs among EBV⁺ B cells in healthy controls (HC) and by disease group and treatment. EBV⁺ B cells from healthy controls were entirely latency 0/I. Latency 0 and latency I are shown as a single category because they are separated only by detection of EBNA1, which is depth-sensitive at these coverages. **b**, Differential expression between EBV⁺ B cells and depth-matched EBV⁻ B cells from the same libraries, plotted as the log₂ fold change against the −log₁₀ P value of a patient-level test. The dotted line marks P = 0.05; colored points are the genes shown in c and the six with the strongest evidence are labeled. **c**, log₂ fold change for thirteen survival-associated genes in EBV⁺ versus depth-matched EBV⁻ B cells, grouped as survival and NF-κB, anti-apoptotic, death-receptor inhibitor and complement inhibitor. Exact P values are given beside each bar, in black where P < 0.05 and in gray otherwise. Because the comparator cells are matched on sequencing depth, these are compositional comparisons: a gene is elevated relative to the rest of the transcriptome in the same cells, and a uniform increase in transcriptional output would not register as a difference. P values are from a one-sample two-sided t-test on per-library log₂ fold changes (n = 11 libraries). **d**, *In vitro* cytotoxicity of the anti-CD20 mAbs rituximab, obinutuzumab and ocrelizumab, the anti-CD19 mAb inebilizumab, CD19-CAR T cells, and the CD19 T-cell engager blinatumomab (CD19 × CD3), against EBV⁻ targets (primary B cells and Ramos cells) and EBV⁺ lymphoblastoid cell lines derived from SLE patients (SLE LCL-1 and SLE LCL-2) across NK cell, CAR T-cell, or T cell effector-to-target ratios. Data are mean ± s.e.m.; differences between EBV⁺ and EBV⁻ targets were assessed by two-way analysis of variance with Sidak’s correction, with exact P values shown at each ratio and ns denoting P ≥ 0.05.

Latency III is the only EBV program in which LMP1 is expressed together with EBNA2 and the EBNA3 proteins.^33^ LMP1 is a constitutively active CD40 mimic that recruits TRAF adaptors and drives both the canonical and the non-canonical NF-κB pathways. EBNA2 drives MYC^28,34^, and NF-κB signaling is required for the survival of EBV-transformed B cells.^35^ Cells carrying this program are activated blasts rather than resting memory B cells. We therefore asked whether the transcriptional state of EBV⁺ B cells in patients matched that expectation. Throughout these analyses EBV⁺ cells were compared with EBV⁻ B cells matched on sequencing depth from the same library. These comparisons are therefore compositional: a gene is elevated relative to the rest of the transcriptome in the same cells, and a uniform increase in total transcriptional output would not register as a difference.

On this basis, all thirteen survival-associated genes examined were expressed at higher levels in EBV⁺ B cells (Fig. 4b, c). Seven reached significance: the TRAF adaptors TRAF1 (*P*=0.019) and TRAF2 (*P*=0.024), the non-canonical NF-κB effector NF-κB2 (*P*=0.024), the canonical subunit RELA (*P*=0.038), the co-chaperone BAG1 (*P*=0.036), the death-receptor inhibitor CFLAR (*P*=0.016) and the complement regulator CD55 (*P*=0.031). The remaining six were also elevated but did not reach significance at this cohort size: REL (*P*=0.203), BCL10 (*P*=0.38), MCL1 (*P*=0.119), BCL2L1 (*P*=0.07), FAS (*P*=0.261) and CD59 (*P*=0.064). Effect sizes were between 1.3- and 2.2-fold (one-sample two-sided t-test on per-library log2 fold changes, n=11 libraries). The uniform direction of this shift is what the latency III program predicts, and because the comparison is depth-matched it reflects enrichment relative to the rest of the transcriptome rather than a general rise in RNA content.

Pre-ranked gene set enrichment analysis of EBV⁺ against depth-matched EBV⁻ B cells from the same post-rituximab libraries identified antigen processing and presentation as the most strongly enriched program (normalized enrichment score 2.05, q = 0.010), together with Toll-like receptor signaling, phagosome, cross-presentation, B-cell receptor signaling and interferon α and γ responses (Extended Data Fig. 5b). Of 155 gene sets reaching a false discovery rate below 0.25 in both the full analysis and a sensitivity analysis restricted to patients contributing at least 5 EBV⁺ cells, 152 were enriched and 3 were depleted; two of the three depleted sets were cell-cycle programs, the G2-M checkpoint (−2.08, q = 0.001) and E2F targets (−1.88, q = 0.004). No gene set changed sign between the two analyses (Extended Data Fig. 5c). EBV⁺ B cells persisting after rituximab therefore occupy a non-proliferative, antigen-presenting state.

### CD19-CAR T cells and a CD19 T-cell engagers eliminate EBV^+^ B cells despite resistance to antibody-mediated cytotoxicity

*In vitro*, three anti-CD20 antibodies (rituximab, ocrelizumab and obinutuzumab) and the anti-CD19 antibody inebilizumab demonstrated reduced cytotoxicity against EBV⁺ lymphoblastoid cell lines relative to EBV⁻ targets at every effector-to-target ratio tested, whereas CD19-CAR T-cells and the CD19 T-cell engager blinatumomab (anti-CD19xCD3) eliminated EBV⁺ and EBV⁻ B-cell targets with comparable efficacy, with no difference at the lowest or highest ratio and a modest reduction in cytotoxicity at the intermediate ratio of 2:1 (Fig. 4d). T cell-mediated killing therefore overcomes a resistance that antibody-mediated cytotoxicity does not: the antibodies were less effective against EBV⁺ than EBV^-^ targets at all three effector-to-target ratios, whereas CD19-CAR T cells and blinatumomab killed EBV⁺ and EBV⁻ targets comparably, differing only at the intermediate 2:1 ratio.

This resistance is unlikely to reflect loss of the target. Inebilizumab showed reduced cytotoxicity against the same antigen that CD19-CAR T cells and blinatumomab engaged effectively (Fig. 4d), indicating that CD19 remains accessible on EBV⁺ targets and that the difference resides in the effector mechanism. Antibody-dependent killing requires complement deposition or Fc receptor-mediated engagement of innate effectors at the surface of the target cell, whereas redirected T-cell killing by CAR T cells and T-cell engagers proceeds through immunological synapse formation and directed release of perforin and granzymes, independent of complement and of Fc receptor engagement.

## Discussion

The ability of CD19-CAR T-cell-mediated ultra-deep B cell depletion to induce sustained, drug-free remission provides a transformative therapeutic paradigm for SLE and other autoimmune diseases^5,6,8,11^. Although CD19-CAR T-cell therapy results in profound depletion of B cells in both blood and tissues^21,36^, the specific pathogenic B cell populations underlying its clinical efficacy have remained undefined.^5,7,8,11^ Here, using EBV-seq, we demonstrate that CD19-CAR T-cell therapy, but not the anti-CD20 monoclonal antibody rituximab, resets the autoreactive EBV⁺ B cell repertoire (Fig. 5). After CD19-CAR T-cell therapy the EBV⁺ compartment carried hallmarks of a naïve state — IgM isotype usage and absence of somatic hypermutation — indicating depletion of the pre-existing EBV⁺ B cell pool and its replacement by newly infected naïve B cells, rather than regeneration from infected memory B cells that survived treatment. Consistent with this, mAbs cloned from EBV⁺ B cells after CD19-CAR T-cell therapy, but not after rituximab, were typically non-autoreactive.

**Fig. 5.**
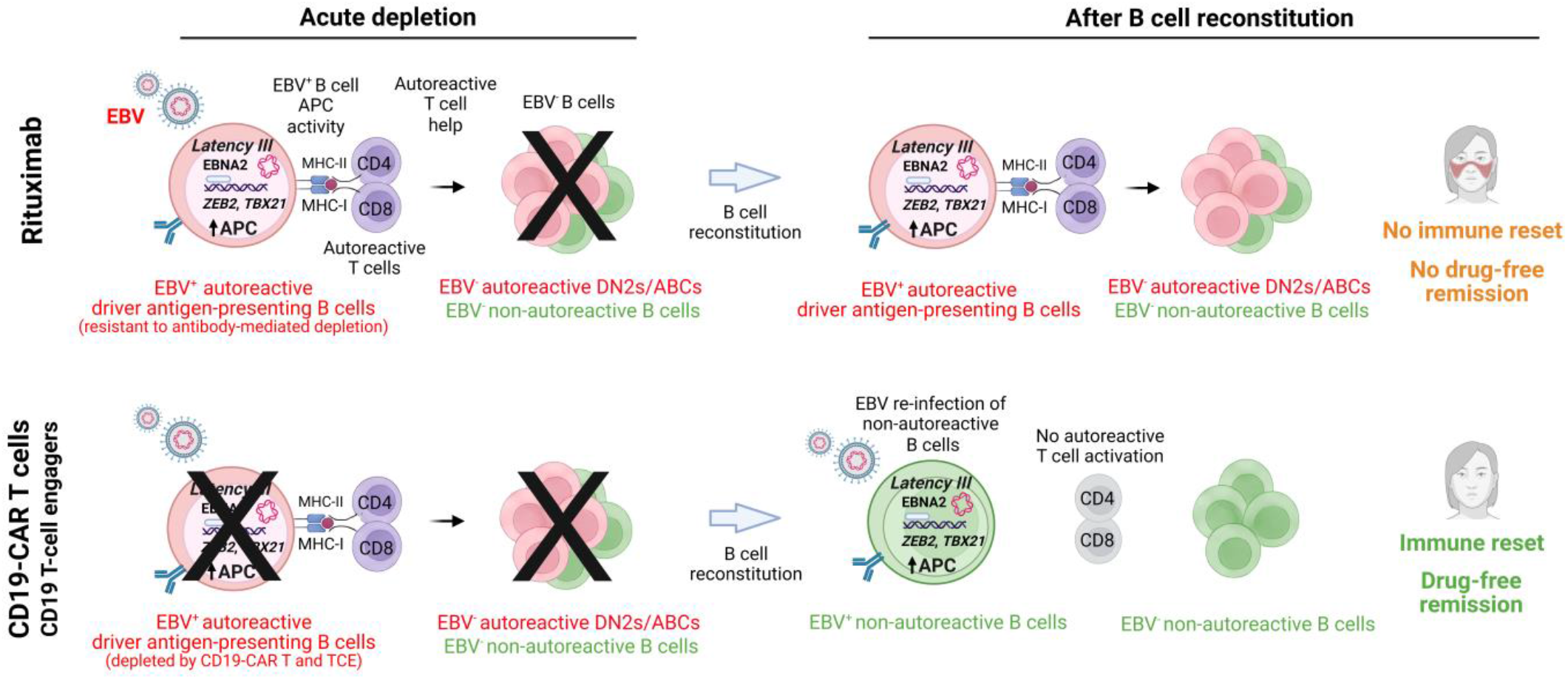
Proposed model for differential resetting of the EBV⁺ autoreactive B cell repertoire by rituximab versus CD19-CAR T cells or CD19 T-cell engagers. Rituximab (top) versus CD19-CAR T cells or CD19 T-cell engagers (bottom), during acute depletion (left) and after B cell reconstitution (right). Before treatment, EBV reprograms autoreactive B cells into driver antigen-presenting cells that carry a latency III program with EBNA2 expression (Fig. 4a) and exhibit elevated MHC class I (MHC-I), MHC class II (MHC-II), antigen-processing, and activation scores relative to EBV⁻ B cells (Fig. 2d–g). These EBV^+^ driver antigen-presenting B cells present autoantigen to autoreactive CD4⁺ and CD8⁺ T cells, and CD4⁺ T cell help in turn activates EBV⁻ autoreactive B cells, including DN2/ABC B cells.^3,4^ **Acute depletion**, Rituximab depletes EBV⁻ autoreactive B cells but not EBV⁺ driver antigen-presenting B cells — consistent with their elevated CD55 and CFLAR (Fig. 4c) and with reduced killing of EBV⁺ targets by anti-CD20 and anti-CD19 antibodies *in vitro* (Fig. 4d). CD19-CAR T cells and T-cell engagers kill independently of complement and Fc receptors and eliminate both EBV⁺ and EBV⁻ targets (Fig. 4d). **After reconstitution**, The surviving EBV⁺ autoreactive driver antigen-presenting B cells continue to present autoantigen and sustain an autoreactive compartment, proposed to preclude immune reset. After CD19-directed T cell killing, the EBV⁺ compartment is repopulated by newly infected naïve IgM B cells (virion) that also exhibit latency III but are non-autoreactive, leaving autoreactive T cells unengaged — proposed to result in immune reset and sustained drug-free remission. Class switching of these non-autoreactive EBV^+^ B cells is expected with time, and EBV may also eventually infect non-autoreactive class-switched cells. If EBV reinfects autoreactive B cells, then autoimmunity can be re-initiated (Fig. 3a, c, d). Autoreactive B cells, pink; non-autoreactive, green; clusters depict the mixed peripheral B cell pool, and depletion by either agent is not antigen-selective. Engaged T cells, purple; unengaged, gray. Black X, depletion; its absence on the EBV⁺ driver B cell denotes resistance. EBNA2-driven induction of *ZEB2* and *TBX21*^3,4^, and the causal link between EBV⁺ repertoire composition and remission, are inferred rather than tested here; the T-cell engager arm rests on *in vitro* data (Fig. 4d). Created in BioRender.

All CD19-CAR T-cell-treated patients achieved initial deep clinical responses and 8 of 12 achieved sustained drug-free remission, in contrast to none of the 15 patients in the rituximab-treated group. Reconstitution shifted the B cell pool as a whole toward unmutated IgM, but the association with sustained remission was confined to the EBV⁺ compartment (Fig. 2b), and after CD19-CAR T-cell therapy the frequency of autoreactivity among EBV⁺ B cells returned to that of the EBV⁻ background (Extended Data Fig. 3c-g). Resetting the EBV^+^ autoreactive B cell repertoire, with concomitant elimination of autoreactive “driver antigen-presenting” B cells^3,4^, may enable peripheral tolerance mechanisms^37,38^ to reassert control. Disease recurred in four of 12 CD19-CAR T-cell-treated patients at months 9, 10, 23 and 30, over a median follow-up for 31 months. The model proposed here accounts for this by two routes: a repertoire reset to a pre-autoimmune state remains susceptible to reinfection of an autoreactive clone by EBV, and where the EBV⁺ compartment is incompletely cleared, pre-existing autoreactive clones persist — as in the patient with SSc (#9), in whom EBV⁺ B cell frequency did not fall after therapy and two EBV⁺ clonotypes were shared across time points (Extended Data Fig. 3a).

Several factors likely underlie the differential ability of CAR T-cells versus rituximab to eliminate EBV^+^ autoreactive B cells and thereby reset the repertoire, including (i) differences in tissue penetration and (ii) distinct mechanisms of cytotoxicity. Rituximab has limited efficacy in depleting tissue-resident B cells^21,36,39^, which likely constrains its ability to eliminate EBV⁺ autoreactive B cells. In contrast, CAR T-cells efficiently traffic into tissues and mediate near-complete depletion of tissue B cells within secondary lymphoid organs and sites of autoimmune inflammation.^21,36^ The second difference is the effector mechanism itself. Anti-CD20 and anti-CD19 antibodies kill through complement deposition and Fc receptor-mediated engagement of innate effectors, both of which act at the target cell surface and both of which the target cell can regulate. EBV⁺ B cells in this cohort upregulated the complement regulator CD55 (Fig. 4c), and complement regulatory proteins are an established determinant of susceptibility to rituximab-mediated lysis in B cell malignancy.^40,41^ CD19-CAR T cells and T-cell engagers bypass this layer entirely, killing through directed granule release independent of complement and of Fc receptor engagement. Tissue access and effector mechanism could each contribute, and the in vitro assays isolate the second by removing tissue access as a variable: against the same targets, the antibody-based agents exhibited reduced cytotoxicity against EBV⁺ cells, whereas the T cell-mediated agents eliminated EBV⁺ and EBV⁻ targets with comparable efficacy. The resistance measured here is therefore a property of the EBV^+^ target cell rather than of drug distribution.

The reset therefore operates on the infected EBV^+^ B cell rather than on the virus or its transcriptional program. The proportion of EBV⁺ B cells in latency III was unchanged from before to after CD19-CAR T-cell therapy, so what distinguishes the post-treatment compartment is not loss of the latency III program but loss of autoreactive specificity among the cells that carry it. B cell-directed therapy does not eliminate the viral reservoir, which resides in epithelial cells^29,33^. EBV genetic variation at EBNA2 has been associated with autoimmune risk loci and, in multiple sclerosis, with disease susceptibility^42^; we did not genotype the virus in this cohort and therefore cannot establish whether the reinfecting strain is that present before treatment. The model predicts that durability depends on whether EBV reinfects an autoreactive clone. Consistent with this, 3 of 7 post-treatment mAbs from the patient with SLE who did not achieve sustained drug-free remission remained HEp-2 reactive, including an unmutated IgM antibody cloned from a latency III B cell (Extended Data Fig. 3b); this observation derives from a single patient and requires confirmation in larger cohorts. Such reinfection of an autoreactive clone is not unexpected: approximately 20% of mature naïve B cells in healthy individuals encode antibodies with antinuclear reactivity^37,43^, a frequency that is substantially increased in autoimmune disease, and autoantibodies are detectable in at-risk individuals years before clinical onset.^44^

The near-universal reinfection of newly generated naïve B cells by 6 months after CD19-CAR T-cell therapy was an unexpected finding. EBV infects both B cells and epithelial cells, with the epithelium serving as a permissive site for viral replication and shedding^29,33^. Moreover, patients with SLE and multiple sclerosis exhibit impaired immune control of EBV^30,31^, which may facilitate lytic reactivation and subsequent reinfection of naïve and/or autoreactive B cells. We hypothesize that the predominant source of reinfection is either the epithelial reservoir or *de novo* exposure through salivary transmission. Re-expansion of EBV⁺ B cells that resisted depletion can also contribute, as shown by the shared EBV⁺ clonotypes in the patient with SSc who relapsed, although no such persistence was detected in any other patient (Extended Data Fig. 3a). Supporting the epithelial reservoir model, ∼50% of SLE patients have detectable EBV DNA in saliva^45^, consistent with ongoing epithelial viral shedding. Further studies will be required to delineate the relative contributions of epithelial reservoirs, residual EBV^+^ B cells, and transmission from another individual to EBV reinfection dynamics.

Here we further show that EBV⁺ B cells in autoimmunity carry the latency III program, which in healthy carriers is confined to newly infected cells and is absent from the circulating memory reservoir.^24,33^ Latency III is the only program in which EBNA2 is expressed, and EBNA2 occupies autoimmune risk loci^28^ and induces expression of ZEB2 and T-bet, which can drive pro-inflammatory autoantigen presentation and thereby systemic autoreactive Th- and EBV⁻ B-cell responses.^3,4^ Together with EBNA1 mimicry of GlialCAM and anoctamin-2^25,26^, and functional mimicry of CD40 by LMP1^27^, this places the EBV⁺ B cell at the convergence of several routes from infection to autoreactivity. Our data identify the EBV^+^ autoreactive B cells as the population whose persistence distinguishes antibody-based depletion from CD19-CAR T-cell therapy.

If the EBV⁺ autoreactive B cell is the population that determines durability, several therapeutic routes follow. The objective is to reproduce *ex vivo* CAR T-cell killing without its manufacturing requirements and with off-the-shelf availability: both *in vivo* CAR T and T-cell engagers provide this potential and have already induced clinical responses in refractory autoimmune disease, although the depth and durability of these responses remain to be fully evaluated.^15–18^ In our *in vitro* assays, a CD19 T-cell engager eliminated EBV⁺ B-cell targets with efficacy comparable to that of CD19-CAR T cells (Fig. 4d). Resetting the EBV⁺ compartment with a T-cell engager therefore appears achievable, but whether the depth of depletion required for repertoire reset can be reached in patients will depend on the dose, exposure duration and tissue penetration achieved clinically. Our data provide a mechanistic rationale for that strategy and specify the EBV⁺ autoreactive B cell population that must be cleared.

Targeting the infected cell or virus directly, rather than the B cell lineage, may also be feasible. Allogeneic EBV-specific T cells are authorized in the European Union for relapsed or refractory EBV-positive post-transplant lymphoproliferative disease after rituximab or another primary therapy^46^, establishing that selective elimination of EBV-infected B cells can be achieved in patients. In an open-label phase I trial, autologous EBV-specific T cells improved clinical outcomes in 7 of 10 patients with progressive multiple sclerosis,^47^ whereas the randomized EMBOLD trial of allogeneic ATA188 did not meet its primary endpoint.^48^ Both studies were conducted in non-active patients with advanced disease and little capacity for clinical improvement.^48^ Vaccination and antivirals targeting EBV provide additional approaches, but their potential impact on established populations of infected autoreactive B cells remains to be characterized.

Several limitations apply: (i) The treatment groups were not randomized and differed in disease composition, center and sample handling, as detailed in Methods, and most comparator-cohort patients contributed a single post-treatment sample; the association between reset depth and sustained drug-free remission is therefore correlative, and with 12 CAR T-cell-treated patients we cannot exclude confounding by disease or cohort. (ii) Analyses were restricted to peripheral blood; the tissue compartments in which EBV⁺ B cells reside were not sampled. (iii) Killing assays used EBV-transformed lymphoblastoid cell lines, which are uniformly latency III and proliferative and may not fully represent EBV⁺ B cells *in vivo*. These assays establish that EBV⁺ B cells are susceptible to T-cell engager-mediated killing, but *in vitro* effector-to-target ratios do not map directly onto clinical exposures and therefore do not predict the dose required to achieve repertoire reset *in vivo.* (iv) Each antibody was tested at a single concentration, so these assays do not establish whether higher anti-CD20 exposure or a CD20 TCE could overcome the resistance. (v) We cannot determine whether depletion of EBV⁺ driver B cells alone is sufficient to induce drug-free remission, or whether concurrent depletion of EBV⁻ autoreactive B cell populations, including DN2/age-associated B cells (ABCs),^49^ is also required. (vi) We did not genotype EBV or sample saliva or epithelial tissue, and therefore cannot determine the source of the reinfecting virus.

In summary, our findings implicate EBV infection of autoreactive B cells, together with latency III-associated, EBNA2-mediated transcriptional reprogramming into activated antigen presenting cells, in the persistence of autoimmune disease. Our data further suggest that therapeutic resetting of the EBV^+^ autoreactive B cell repertoire to a non-autoreactive naïve state may underlie sustained drug-free remission. These observations provide a mechanistic rationale for the development of EBV-targeted therapeutic strategies for autoimmune diseases.

## Methods

### Patients

Three cohorts were analyzed. Twelve patients with treatment-refractory autoimmune disease (SLE, n=6; IIM, n=3; SSc, n=3) were enrolled at Universitätsklinikum Erlangen in the CASTLE basket trial (NCT06347718)³ and treated with a single infusion of CD19-CAR T cells (Zorpo-cel [MB-CART19.1, Miltenyi Biomedicine] or KYV-101 [Kyverna]); peripheral blood was collected before and 6 months after infusion, except in two patients with SLE for whom only the 6-month sample was available. Seven patients with SLE were enrolled at Leiden University Medical Center in the Synbiose-II trial (NCT0^33^47159) and treated with rituximab plus belimumab; peripheral blood was collected at week 0 and week 44. Eight patients with SLE treated with rituximab at Stanford University contributed a single sample collected approximately 6 months after infusion. Eight EBV-seropositive healthy donors, drawn from our previously described cohort (PRJNA1301449), served as comparators. All patients provided written informed consent under protocols approved by the institutional review boards of Universitätsklinikum Erlangen, Leiden University Medical Center and Stanford University. Patient characteristics are given in Table 1 and Supplementary Tables 1 to 3.

Sample preparation differed between the cohorts, which constrains which comparisons are valid. The rituximab (Synbiose-II) samples were all pan-B enriched by negative selection (StemCell EasySep), whereas the CD19 CAR-T cohort comprised approximately half whole PBMC and half enriched B cells, with the sort fraction differing between the pre- and post-treatment sample of the same patient in some cases; the measured B-lineage fraction was 80.6% across the rituximab cohort against 38.2% across the CAR-T cohort. All frequencies reported here are therefore proportions within the B compartment, or EBV+ cells per 10,000 B cells, both of which are invariant to a shifting non-B fraction. No pre- to post-treatment comparison was computed outside the B compartment, and the two cohorts were not pooled for any per-library uniformity test.

### EBV-seq and Bioinformatic Analysis

For the studies reported here, we used EBV-seq.2, which uses refined versions of our previously reported EBV-seq primers, library preparation methods, and informatics described by Younis et al.^3^. EBV-seq.2 uses the 10x Genomics 5′ single-cell RNA-seq platform with capture of EBV viral RNA during reverse transcription and library amplification. The refined 16-primer panel, spanning 13 genes, targets the most informative EBV genes for latency phase classification and lytic activity, with primers selected for alignment specificity against both type 1 and type 2 genomes, screened against the human genome to exclude 3’-anchored mispriming, and with concentrations weighted to target abundance. Of the 53 libraries generated for this study, 41 used the EBV-seq.2 panel and the remaining 12 used the previously published panel^3^. Because each captured molecule extends from the transcript cap to the gene-specific primer, amplicon length is set by the target, and library preparation was restructured so that both long and short amplicons are represented. Short amplicons, including the EBERs, LMP1 and the lytic targets, are carried in a dedicated library prepared without fragmentation and amplified selectively, so that untargeted transcriptome cDNA is not amplified. The informatics were refined as described below. Reads were aligned against a combined GRCh38 and EBV reference carrying a curated EBV annotation, and then subset to the EBV contig; host sequence is retained in the index because EBV has genuine host homology and host reads otherwise mismap onto the 172 kb viral genome.

#### Latency panel

Calling was performed on a fixed panel of EBV latency genes: EBNA2 (BYRF1), EBNA1, EBNA-LP, EBNA3A, EBNA3B, EBNA3C, LMP1, LMP2, EBER1, EBER2 and the BART transcripts. Calling was performed on annotation-level counts over a fixed panel of EBV latency genes, which is not restricted to the captured targets.

#### Ambient reference

Calling rests on a single principle: in droplet single-cell data most EBV reads are ambient, so the presence of EBV reads does not by itself identify an infected cell. For each sample the ambient EBV profile was estimated from cells that cannot harbor EBV infection, preferring non-B cells from the same library and falling back to empty droplets where fewer than 50 non-B cells were available. Samples with fewer than 100 empty droplets used the pooled cohort ambient profile. Gene proportions were estimated with a pseudocount of 1.

#### Depth model

The expected ambient latency-gene UMI count for each cell was predicted from its sequencing depth by a generalized linear model fitted on the ambient reference cells of the same sample, requiring at least 50 non-B cells. Where a sample had fewer, the model was fitted instead on its B cells, trimmed at the 95th percentile of the response to prevent genuinely infected cells from inflating the ambient expectation.

#### Test statistic

Each cell was scored by two deviance components against its own predicted ambient expectation. An enrichment deviance tested whether the total latency UMI count exceeded expectation, D_enrich = 2[y·log(y/λ) − (y − λ)], where y is the observed and λ the predicted count. A composition deviance tested whether the distribution of counts across panel genes departed from the ambient gene proportions, D_comp = 2·Σ_g y_g·log[(y_g/y)/p_g]. The two were combined and referred to a null distribution obtained by Monte Carlo simulation (100,000 draws) at the cell’s predicted ambient rate, with cells binned geometrically so that the binned and exact rates differ by no more than 10%, and with analytic negative-binomial tail extrapolation beyond the resolution of the simulation. Overdispersion was estimated per sample and constrained to between 1 and 10.

#### Calling rule

P values were corrected by the Benjamini-Hochberg procedure stratified on the predicted-ambient bin, and a cell was called EBV⁺ at a false discovery rate below 0.01, subject to a floor of at least 2 latency UMIs and an observed count not below the ambient expectation, so that a cell cannot be called on gene composition alone.

#### Reproducibility

All stochastic steps used a fixed random seed. Splice-junction evidence spanning the EBV W repeat was used to corroborate calls.

Latency programs were assigned to each called cell from the detection of the defining viral genes: latency III, EBNA2 detected together with EBNA3B, EBNA3C or LMP1; latency II, EBNA2 absent with LMP1 detected; latency I, EBNA2 and LMP1 absent with EBNA1 detected; latency 0, EBNA2, LMP1 and EBNA1 absent with EBER1 or EBER2 detected. Cells additionally expressing the immediate-early lytic genes BZLF1 or BRLF1 were annotated as lytic, and cells meeting none of these criteria were left unassigned. Only latency I, II and III are used in the analyses reported here. Because assigning latency III requires detecting EBNA2, latency assignment is sensitive to sequencing depth; comparisons between latency classes were therefore made against depth-matched backgrounds as described below, and per-class cell counts are reported alongside every such comparison.

### B-cell subset classification

B cells were assigned to subsets using CITE-seq surface protein together with RNA isotype expression. Antibody-derived tag (ADT) counts were centered-log-ratio (CLR) normalized per cell before thresholding. Subsets were defined on CD27, IgD, CD21, CD11c and a CD138 plasmablast score, with switched versus unswitched identity resolved by the ratio of IgG and IgA to IgM and IgD transcripts: naïve (CD27^-^IgD^+^), unswitched memory (CD27^+^IgD^+^), switched memory (CD27^+^IgD^-^), CD27^+^CD21^low^ memory, DN1 (CD27^-^IgD^-^CD11c^-^), DN2/ABC (CD27^-^IgD^-^CD11c^+^) and plasmablasts.

### Monoclonal antibody expression

Paired heavy- and light-chain variable region sequences were identified from single EBV^+^ B cells by single-cell V(D)J sequencing. Variable regions were cloned as human IgG1 (G1m3 allotype) with matched kappa or lambda light chains, and the recombinant antibodies were expressed and purified by GenScript Biotech. Monoclonal antibodies from EBV-negative B cells of the same patients, used as the repertoire background in Extended Data Fig. 3, were cloned and expressed by the same procedure.

### HEp-2 indirect immunofluorescence and ELISA for anti-nuclear antibody reactivity

Autoreactivity was assessed using commercial HEp-2 indirect immunofluorescence assays. Purified mAbs were diluted to 1–10 μg/mL and applied to HEp-2 substrate slides according to the manufacturer’s protocol. Bound antibodies were detected using an FITC-conjugated anti-human IgG secondary antibody. Staining patterns were classified by blinded evaluation as nuclear homogeneous, nuclear speckled, cytoplasmic, nucleolar, or negative.

### LCL generation

Spontaneous LCLs (sLCL) were generated from SLE patient PBMCs using a modified standard protocol^50^. T cells were depleted from PBMCs by magnetic-activated cell sorting (MACS) using CD3 MicroBeads (Cat#130-050-101; Miltenyi Biotec). The resulting cells were cultured in RPMI-1640 supplemented with 20% FBS, 1% glutamine, and 1% penicillin/streptomycin, with cyclosporin A (1 μg/mL) added to suppress residual T cell activity. Cultures were maintained by weekly refeeding with fresh complete media for 6–8 weeks, until actively proliferating lymphoblastoid cell clusters emerged. These sLCL clusters were subsequently harvested, expanded for an additional 2–4 weeks, and cryopreserved for downstream analyses.

### *In vitro* cytotoxicity assays

Healthy donor B cells were isolated by negative selection using the EasySep™ Human B Cell Isolation Kit (Cat#17954; STEMCELL Technologies). The Ramos (RA 1) cell line was obtained from American Type Culture Collection (ATCC; CRL-1596). For NK cell–mediated cytotoxicity assays, untouched NK cells were isolated from healthy donor PBMCs using the EasySep™ Human NK Cell Isolation Kit (Cat#17955; STEMCELL Technologies) and pre-activated overnight in complete medium (RPMI-1640 supplemented with 10% FBS, 1% glutamine, and 1% penicillin/streptomycin) in the presence of IL-2 (10 ng/mL; Cat#202-IL, R&D Systems). Activated NK cells (effectors) were then co-cultured with target cells (primary B cells, Ramos cells, or sLCLs) at effector-to-target (E:T) ratios of 1:1, 2:1, and 5:1, in the presence of rituximab biosimilar (10 μg/mL; Cat#SIM0008, Bio X Cell), obinutuzumab (10 μg/mL; Cat#A2023, Selleck Chemicals), ocrelizumab (10 μg/mL; Cat#HY-P9960, MedChemExpress), or inebilizumab (10 μg/mL; Cat#HY-P99113A, MedChemExpress). For T-cell engager cytotoxicity assays, T cells were isolated by negative selection using the EasySep™ Human T Cell Isolation Kit (Cat#17951; STEMCELL Technologies) and co-cultured with target cells at E:T ratios of 1:1, 2:1, and 5:1 for 24 hours in the presence of blinatumomab biosimilar (100 ng/mL; Cat#ICH5025, IchorBio). For CD19 CAR T cell–mediated cytotoxicity, anti-CD19 CAR T cells (eGFP) were obtained from BPS Bioscience (Cat#78789) and co-cultured with target cells at E:T ratios of 1:1, 2:1, and 5:1 for 24 hours. After 24 hours, cells were harvested and analyzed for target cell lysis by flow cytometry using the Zombie Violet™ Fixable Viability Kit (Cat#423113; BioLegend), according to the manufacturer’s protocol. Gating strategy detailed in Supplementary Fig. 3.

### Statistics and reproducibility

#### Sample size

No statistical method was used to predetermine sample size. All patients meeting enrollment criteria available samples were included: CD19-CAR T-cell-treated SLE (n=6; 4 with paired pre- and post-treatment samples and 2 with post-treatment samples only), IIM (n=3, all paired) and SSc (n=3, all paired), and rituximab-treated SLE (n=7 with paired week 0 and week 44 samples; paired comparisons were restricted to this arm, with the remaining 8 patients contributing post-treatment samples only). Cohort sizes reflect the availability of these rare treatment cases rather than a power calculation, and the study is correspondingly powered to detect only large, consistent effects.

#### Replication

Each clinical sample was sequenced once; independent biological replication is not possible for material of this kind. Reproducibility was therefore assessed computationally, and the checks applied depended on the type of finding. Transcriptional findings from the EBV^+^ versus EBV^-^ comparison were required to be independent of the number of EBV^+^ cells a patient contributed (Spearman correlation between per-patient effect size and cell count), to hold in the better-powered half of patients, and to exceed size-matched random gene sets. Candidate transcriptional findings that did not meet all of these criteria are not reported. Compositional findings were additionally tested at two definitions of the classifying threshold to confirm they were not an artifact of where the cut was placed.

#### Randomization

This was an observational study of patients receiving clinically indicated therapy. Allocation to CD19-CAR T-cell or rituximab treatment was not randomized and randomization was not applicable. Comparisons were paired within patient wherever possible, so that each patient served as their own control.

#### Blinding

Blinding to treatment group was not possible for sample acquisition or sequencing, as group membership is intrinsic to the clinical samples. Computational assignment of EBV status, B-cell lineage and B-cell subtype was performed by algorithms blind to treatment group and timepoint.

#### Statistical tests

All tests were two-sided. Paired pre- and post-treatment comparisons used the Wilcoxon signed-rank test; unpaired group comparisons used the Mann-Whitney U test; categorical comparisons used Fisher’s exact test. Paired analyses were restricted to patients contributing both pre- and post-treatment samples and therefore exclude the two patients with SLE and the eight rituximab-treated patients who contributed a single post-treatment sample. For single-cell comparisons, values were aggregated to one measurement per patient, or per library for the depth-matched pseudobulk comparisons in Fig. 4b-c. EBV^+^ cells were compared with EBV^-^ B cells matched on sequencing depth from the same library, with immunoglobulin and mitochondrial genes excluded from the library size. Pre-ranked gene set enrichment analysis was performed with gseapy on genes ranked by a moderated paired t statistic across patients, against the KEGG, Reactome and MSigDB Hallmark collections, with Benjamini-Hochberg correction within each collection. Robustness was assessed by repeating the analysis restricted to patients contributing at least 5 EBV⁺ cells and, separately, to genes detected in all patients; sets reported as robust reached a false discovery rate below 0.25 in both passes with no change in sign. Multiple comparisons were corrected by the Benjamini-Hochberg procedure and are reported as q or FDR. With n=7 paired patients the smallest attainable Wilcoxon signed-rank P value is 0.0156, which indicates that all patients agreed in direction rather than that the effect was large; P values at this floor are interpreted accordingly.

#### Software

Alignment and quantification used Cell Ranger with the GRCh38-2024-A transcriptome and the Ensembl 7.1.0 VDJ reference. B-cell lineage was assigned by the consensus of CellTypist v1.6.3 (Immune_All_Low, majority voting over clusters) and SingleR v2.12.0 against the Monaco immune reference (celldex v1.20.0), both applied at cluster level; doublets were identified with scDblFinder v1.24.10. Clustering and normalization used Seurat v5.5.1, scran v1.38.1, bluster v1.20.0 and scanpy v1.10.4. Gene set enrichment used preranked GSEA (gseapy). Analyses ran in R v4.5.2 with Bioconductor v3.22, and in Python (v3.6.8 for cluster pseudobulk, v3.9 for figure generation).

## Acknowledgments

We thank the Robinson lab members for scientific discussions.

## Author contributions

Y.C. and M.P. contributed equally to this work. Y.C. designed the primer panel used to generate 41 of the 53 single-cell libraries analyzed in this study, comprising all Synbiose-II, all rituximab-treated and all IIM samples and a subset of SLE CD19-CAR T-cell samples. Y.C. and L.S.v.D. designed the single-cell experimental approach for the Synbiose-II arm. Y.C. performed the computational analyses, including EBV calling, B-cell subset classification, differential gene expression analysis and figure generation, and wrote the manuscript. Y.C., M.P. and L.S.v.D. performed single-cell library preparation and sequencing. M.P. generated the lymphoblastoid cell lines, prepared the remaining 12 single-cell libraries (the SSc and remaining SLE CD19-CAR T-cell samples) using the published EBV-seq primer panel, and performed the monoclonal antibody, T-cell engager and CD19-CAR T-cell cytotoxicity experiments. L.S.v.D., E.J.A., C.v.K. and Y.K.O.T. designed and conducted the Synbiose-II trial and provided clinical data and samples. S.Y. designed the CITE-seq antibody panel, developed the EBV-seq method, provided the primer oligonucleotides used for those 12 libraries, and advised on application. S.R. performed the HEp-2 indirect immunofluorescence staining and ELISAs. M.Z. contributed to development of the EBV calling approach and independently implemented and tested it. Y.Ch. enrolled and clinically characterized the Stanford rituximab-treated patients. S.P. and T.V.L. maintain the rheumatology biobank and provided patient samples. M.H., A.W., F.M., A.M. and A.B. enrolled and clinically characterized the CD19-CAR T-cell-treated patients and provided samples. R.G.-B., G.S. and W.H.R. jointly supervised the work. W.H.R. conceived the study and obtained funding. All authors reviewed and approved the final manuscript.

## Data availability

Raw single-cell sequencing data generated in this study will be deposited before publication. Sequencing data from the Stanford samples will be deposited in the NCBI Sequence Read Archive. Sequencing data from the Erlangen CD19-CAR T-cell cohort and from the Leiden Synbiose-II cohort will be deposited in the European Genome-phenome Archive, where access is controlled and governed by the consent under which those samples were obtained and by the applicable European data protection requirements; requests will be reviewed by the data access committee of the originating institution. Accession numbers will be provided at proof stage. Single-cell sequencing data from the healthy control donors were reported previously and are available under BioProject PRJNA1301449.

Processed data sufficient to reproduce every figure will be provided as Source Data and comprise: per-cell EBV call tables including per-gene latency UMI counts, the deviance statistic, the false discovery rate and the latency assignment for every cell tested; per-cell B-cell subset assignments with the normalized surface protein values used to derive them; and the per-panel summary tables underlying each main and supplementary figure.

## Materials availability

The spontaneous lymphoblastoid cell lines generated in this study were derived from the peripheral blood of patients enrolled in clinical trials and are not available for distribution, because the consent under which the source samples were obtained does not extend to transfer of patient-derived cell lines to third parties. The protocol used to generate equivalent lines from primary B cells is described in Methods and ref. 59. Recombinant monoclonal antibodies cloned from single patient-derived B cells are available from the corresponding author under a materials transfer agreement, and the corresponding heavy- and light-chain variable region sequences are included in the deposited V(D)J data. All other reagents are commercially available and are identified in Methods.

## Code availability

No general-purpose software package is released with this study. EBV calling, latency assignment, B-cell subset classification and the depth-matched pseudobulk comparison were performed with an in-house implementation written for this work. That implementation is specified in Methods at a level of detail sufficient for independent reimplementation, including the latency gene panel, the ambient model, the test statistics and every threshold and random seed. The implementation is available from the corresponding author on reasonable request. All other analyses used the publicly available software listed under Software, applied as described in Methods, and the deposited per-cell tables allow every reported figure to be regenerated without repeating the upstream alignment and calling steps.

## Extended data figures

**Extended Data Fig. 1.**
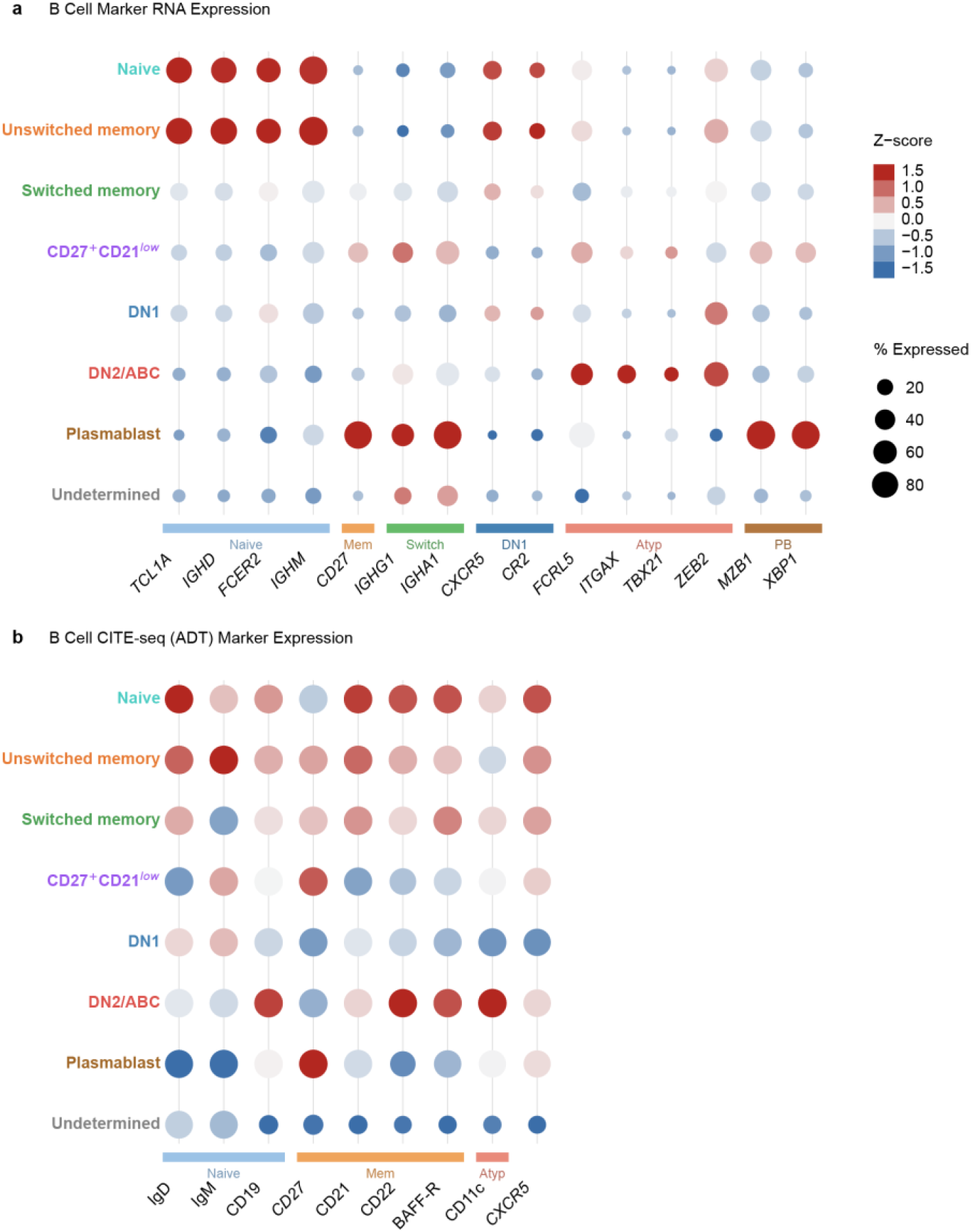
Validation of B-cell populations by RNA and surface protein expression. Dot plots show canonical marker expression across the eight B-cell populations defined in this study, including the cells that could not be assigned to a subset (Undetermined). Dot size indicates the percentage of cells with detectable expression and color indicates the z-score of mean expression across populations (clipped at ±1.5). Marker groupings are indicated by colored bars beneath each panel. **a**, RNA expression of B-cell subtype markers. **b**, Surface protein expression measured by CITE-seq (antibody-derived tags, ADT) in the CD19-CAR T-cell-treated cohorts. Samples obtained after rituximab treatment were processed with a separate CITE-seq panel and are not included in b. Cluster annotations are supported by concordant RNA and ADT expression, including elevated ZEB2 with a CD27⁺CD21ˡᵒʷ surface phenotype, co-expression of ITGAX, TBX21, ZEB2 and FCRL5 in DN2/age-associated B cells, and expression of MZB1 and XBP1 in plasmablasts.

**Extended Data Fig. 2.**
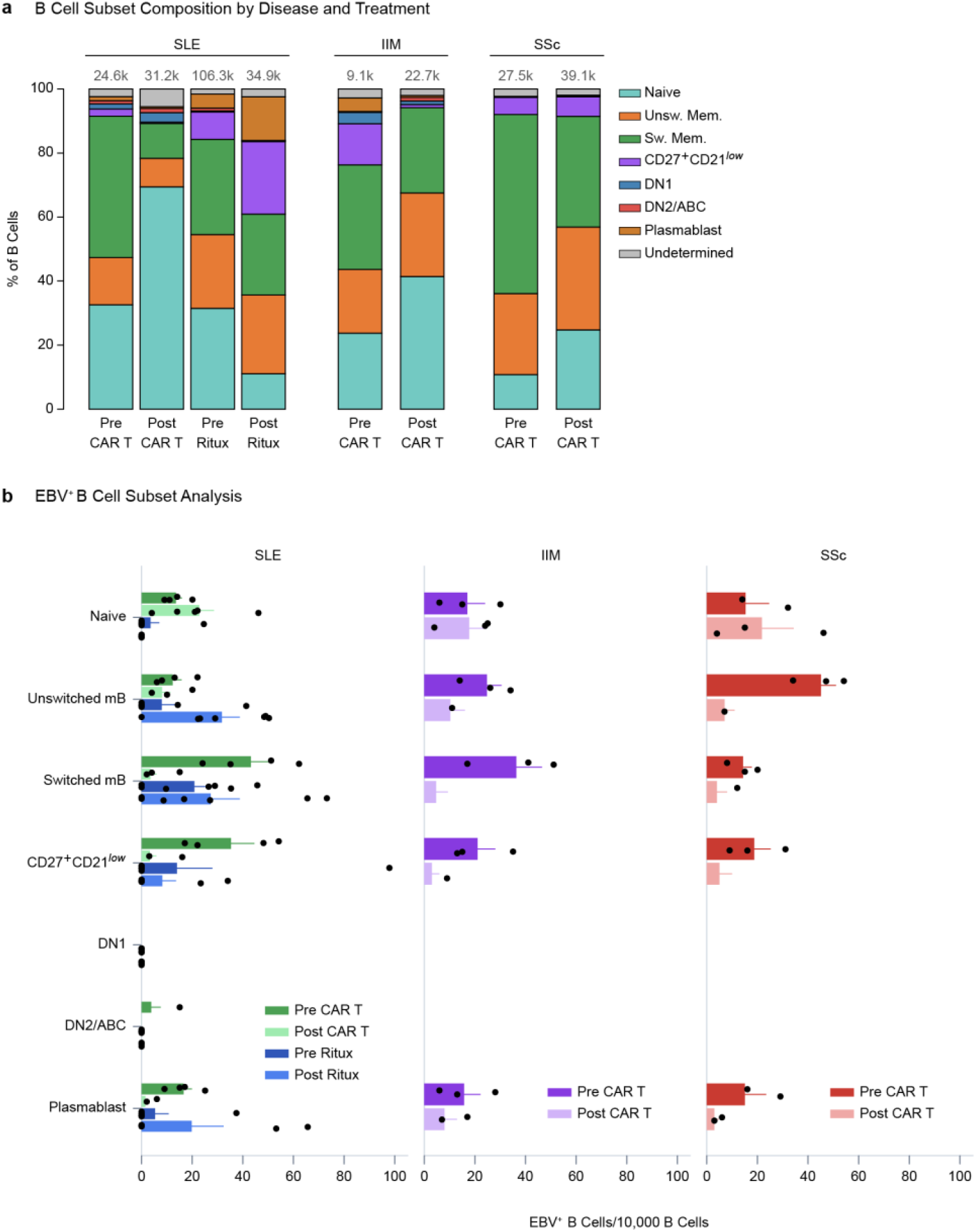
B-cell subset composition and distribution of EBV⁺ B cells according to disease and treatment. **a**, Composition of B-cell subsets, shown as stacked bars giving the proportion of each subset as a percentage of total B cells for each disease group and treatment condition. The number of B cells contributing to each bar is given above it. **b**, Distribution of EBV⁺ B cells across B-cell subsets, expressed as the number of EBV⁺ B cells per 10,000 B cells within each subset, shown separately for SLE, IIM and SSc. Bars indicate cohort means and horizontal lines indicate 95% confidence intervals (Wilson method); dots represent individual patients. Colors denote treatment conditions within each disease group.

**Extended Data Fig. 3.**
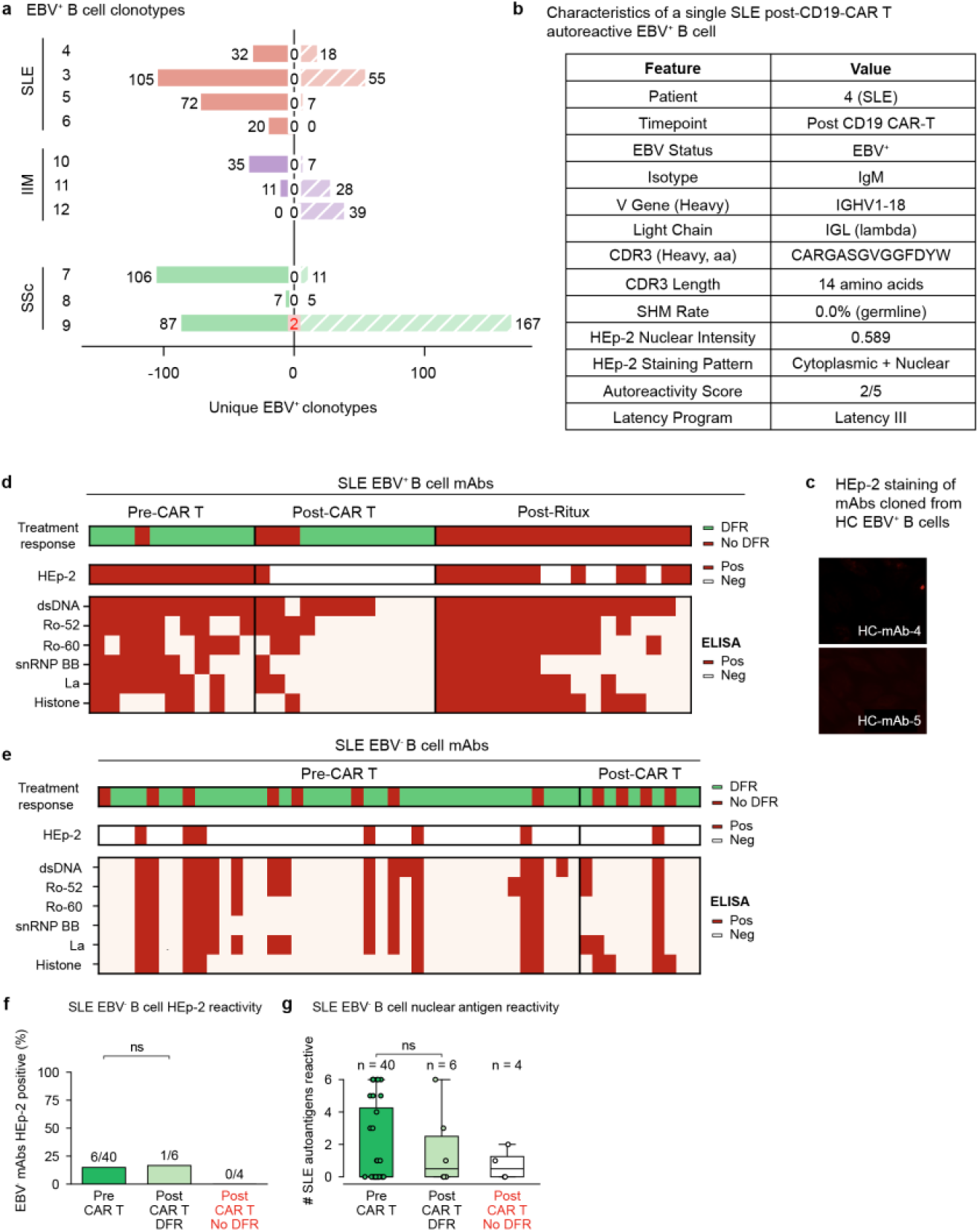
Clonal analysis of EBV⁺ B cells and autoreactivity of monoclonal antibodies from EBV⁺ and EBV⁻ B cells. **a**, Clonal overlap of EBV⁺ B cells before and after CD19-CAR T-cell therapy for each patient with paired samples. Diverging bars indicate clonotypes unique to each time point and the center value indicates clonotypes shared between time points. Clonotypes were defined on immunoglobulin heavy-chain (IGH) sequences obtained with Cell Ranger V(D)J. Two shared clonotypes were detected in SSc patient 9 and none in any other patient. **b**, Features of mAb-3, cloned from an EBV⁺ B cell obtained after CD19-CAR T-cell therapy from SLE patient 4, the one patient with SLE who relapsed. **c**, HEp-2 staining of mAbs cloned from EBV⁺ B cells of healthy controls. **d**, HEp-2 reactivity, treatment response and ELISA reactivity against six SLE-associated nuclear antigens for every mAb cloned from an EBV⁺ B cell, grouped by treatment context. **e**, The same measures for every mAb cloned from an EBV⁻ B cell of the same patients. **f**, Percentage of HEp-2-positive mAbs from EBV⁻ B cells by treatment group; two-sided Fisher’s exact test. **g**, Number of SLE-associated nuclear autoantigens recognized per EBV⁻ B-cell-derived mAb; n indicates the number of mAbs per group, each dot is one mAb, horizontal lines within the boxes indicate medians and the boxes indicate interquartile ranges; two-sided Mann–Whitney U test. In **f** and **g**, red labels denote patients not in sustained drug-free remission.

**Extended Data Fig. 4.**
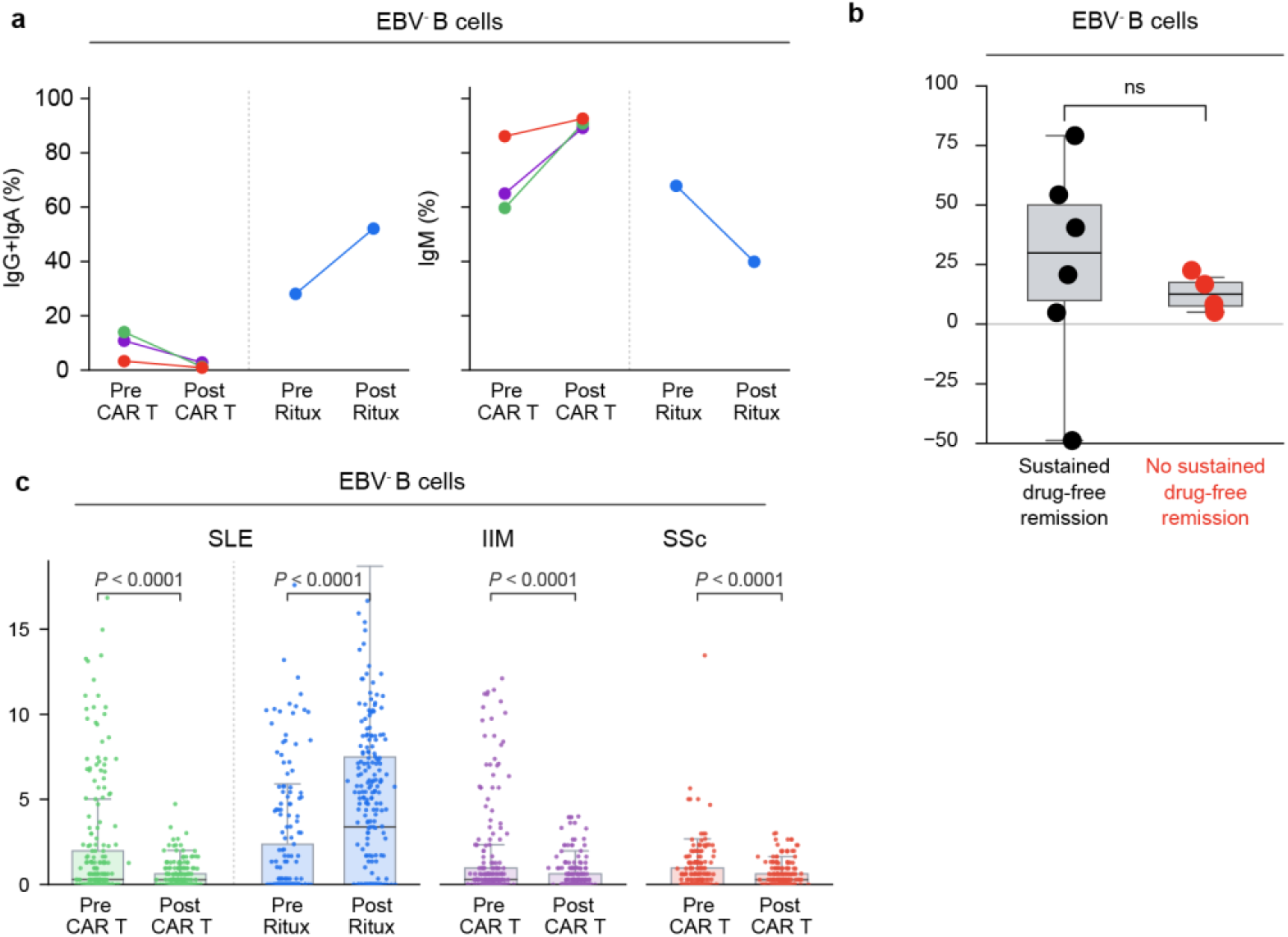
Isotype usage and somatic hypermutation in EBV⁻ B cells. **a**, Percentage of class-switched (IgG⁺ or IgA⁺, left) and IgM⁺ (right) EBV⁻ B cells before and after CD19-CAR T-cell therapy in SLE, IIM and SSc, and before and after rituximab in SLE, plotted as in Fig. 2a. **b**, V-region somatic hypermutation rate of EBV⁻ B cells by disease group and treatment, plotted as in Fig. 2c; each dot represents one cell and bars show the median and interquartile range. These panels provide the EBV⁻ comparator for Fig. 2a, c: the shift toward unmutated IgM after CD19-CAR T-cell therapy is seen in both compartments, consistent with reconstitution of the whole B-cell pool from naïve precursors, whereas somatic hypermutation rises after rituximab in the EBV⁻ compartment.

**Extended Data Fig. 5.**
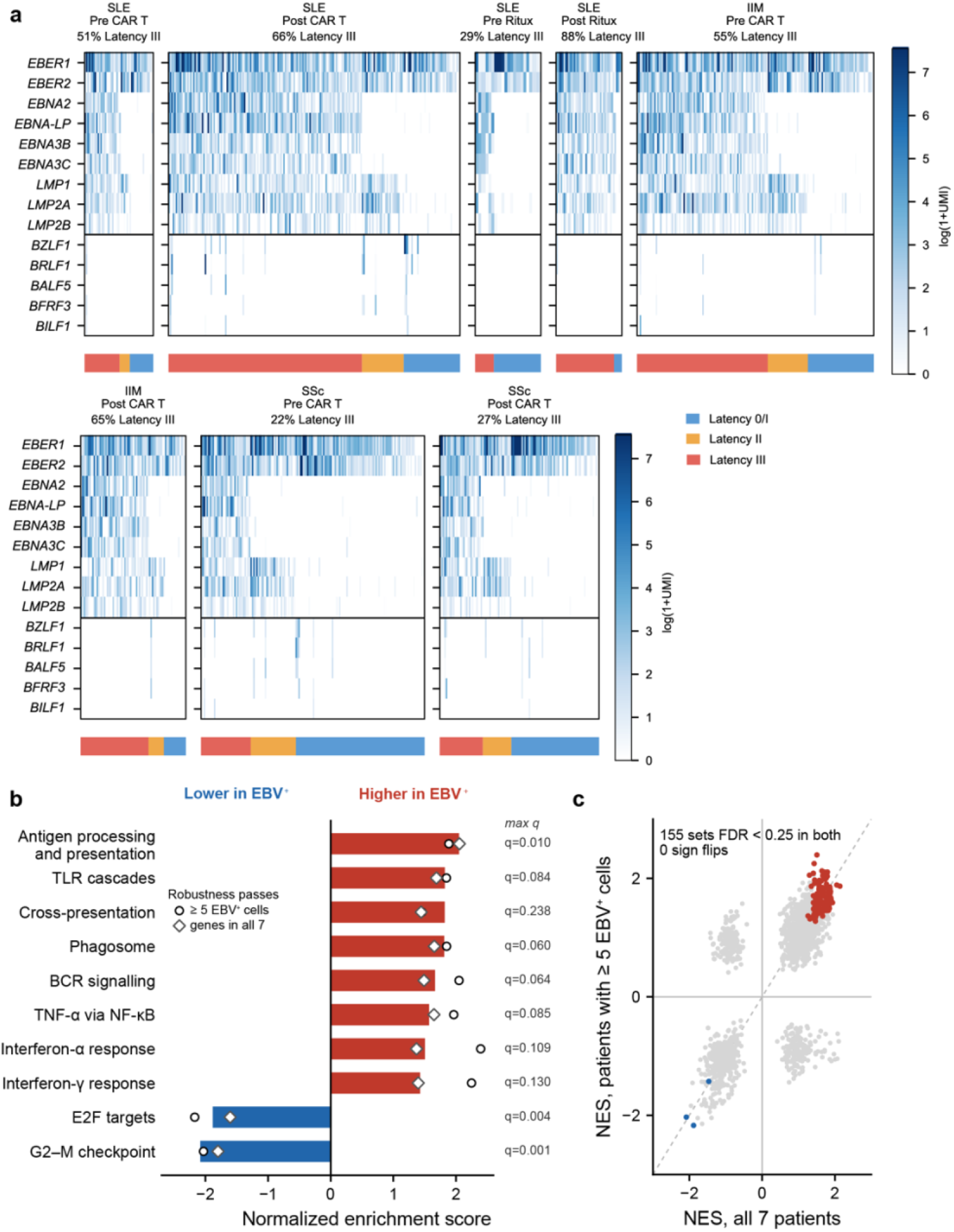
EBV latency programs across disease groups and treatments, and pathway enrichment in EBV⁺ B cells after rituximab. a, EBV gene expression in individual EBV⁺ B cells, shown as log(1+UMI), by disease group and treatment: SLE before (n = 94 cells) and after (n = 399) CD19-CAR T-cell therapy; SLE before (n = 21) and after (n = 51) rituximab; IIM before (n = 324) and after (n = 144) CD19-CAR T-cell therapy; and SSc before (n = 306) and after (n = 218) CD19-CAR T-cell therapy. Each column is one cell. Latency-associated genes are shown above the horizontal rule and lytic-cycle genes below it. The bar beneath each panel gives the latency programme assigned to each cell (latency 0/I, blue; latency II, orange; latency III, red), and cells are ordered by latency programme within each panel. The percentage of cells in latency III is given above each panel. Healthy controls are not shown; all healthy-control EBV⁺ B cells were in latency 0/I (Fig. 4a). b, c, Pre-ranked gene set enrichment analysis of EBV⁺ versus depth-matched EBV⁻ B cells from post-rituximab samples of 7 patients with SLE in the Synbiose-II trial (44 EBV⁺ cells; each EBV⁺ cell matched to its 20 nearest EBV⁻ B cells by total UMI within the same library; immunoglobulin and mitochondrial genes excluded from the ranking). Genes were ranked by a moderated paired t statistic computed across patients, so the patient, not the cell, is the unit of replication. **b**, Normalized enrichment scores for selected gene sets, colored by direction: red, higher in EBV⁺ cells; blue, lower. Bars give the normalized enrichment score across all 7 patients. Overlaid markers give the value from each robustness pass: circles, patients contributing at least 5 EBV⁺ cells; diamonds, restricted to genes detected in all 7 patients. The value beside each bar is the larger of the two false discovery rates across passes. **c**, Normalized enrichment score across all 7 patients plotted against the value from the pass restricted to patients contributing at least 5 EBV⁺ cells, for all 1,733 gene sets tested. Colored points are the 155 sets reaching a false discovery rate below 0.25 in both passes; all 155 agree in direction and none changes sign. Gene sets are from KEGG, Reactome and MSigDB Hallmark. False discovery rates are Benjamini-Hochberg corrected within each collection.

