## Supplemental Material for "CD19-CAR T cells reset the EBV^+^ autoreactive B cell repertoire in autoimmune disease"

### Supplementary Material

Supplementary Figure 1. Additional HEp-2 indirect immunofluorescence staining of monoclonal antibodies derived from $EBV^+$ B cells.

Supplementary Figure 2. HEp-2 indirect immunofluorescence staining of monoclonal antibodies derived from $EBV^-$ B cells.

Supplementary Table 1. Characteristics of 12 Patients Treated with CD19-CAR T Cells.

Supplementary Table 2. Characteristics of SLE Patients Treated with Rituximab.

Supplementary Table 3. Individual patient characteristics and clinical response of the 7 patients with SLE treated with rituximab plus belimumab in the Synbiose-II trial.

*Supplementary References*

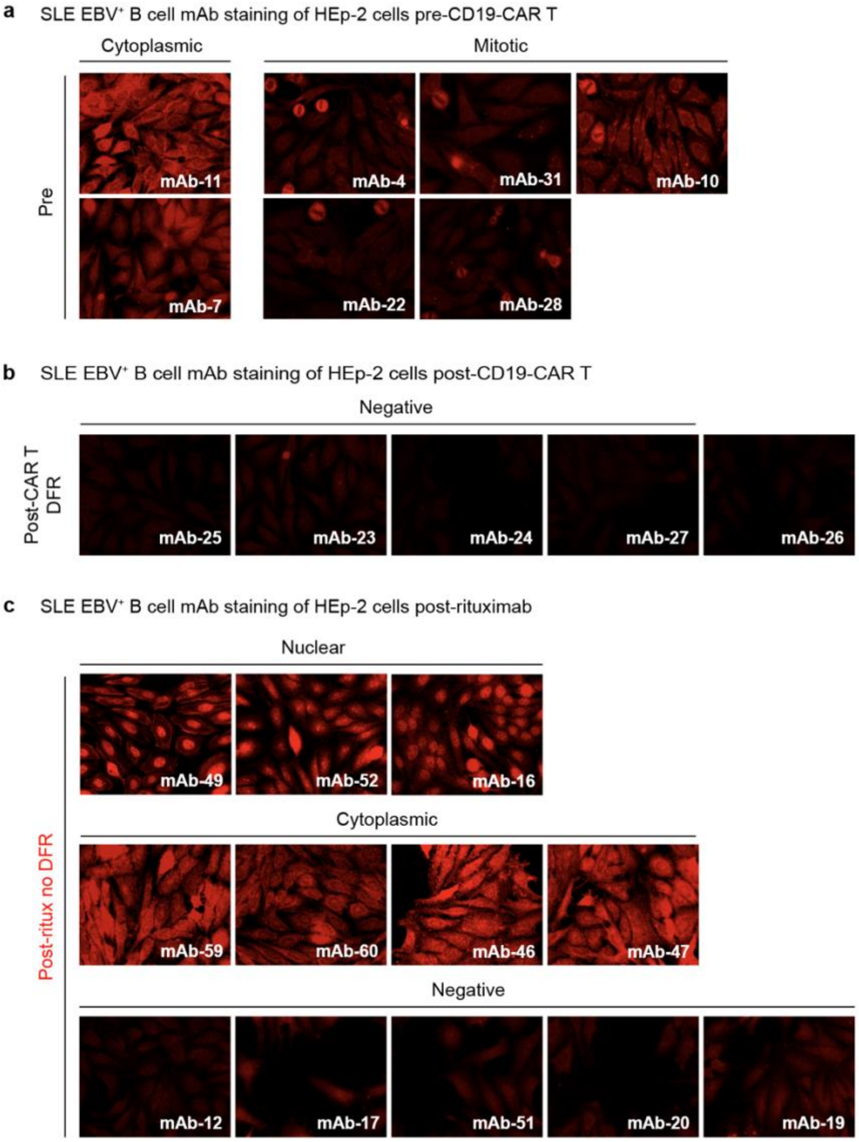

**Supplementary Figure 1. Additional HEp-2 indirect immunofluorescence staining of monoclonal antibodies derived from EBV$^{+}$ B cells.**

Staining of the mAbs cloned from EBV$^{+}$ B cells that are not shown in Fig. 3. **a**, mAbs obtained before CD19-CAR T-cell therapy, grouped by staining pattern. **b**, mAbs obtained after CD19-CAR T-cell therapy from patients in sustained drug-free remission (DFR); all were HEp-2 negative. **c**, mAbs obtained after rituximab treatment, grouped by staining pattern; none of these patients were in DFR. Images were acquired at 20× magnification on HEp-2 substrate slides at an antibody concentration of 10 µg per millilitre. [AUTHOR CHECK: confirm that Fig. 3 and this figure together account for all 14, 10, 7 and 19 mAbs quantified in Fig. 3c, and add scale bars.]

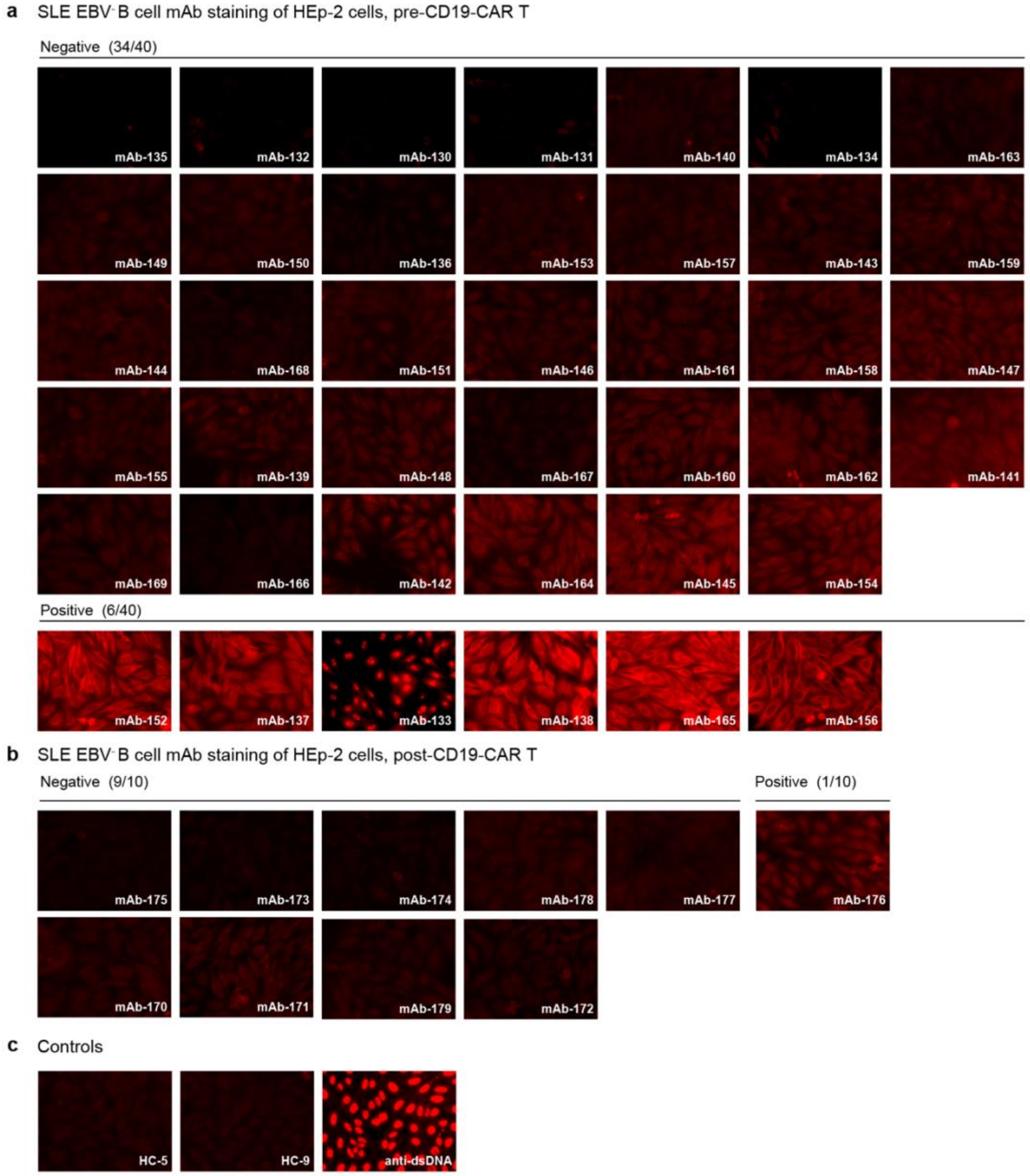

**Supplementary Figure 2. HEp-2 indirect immunofluorescence staining of monoclonal antibodies derived from EBV⁻ B cells.**

**a**, Staining of all mAbs cloned from $EBV^-$ B cells obtained before CD19-CAR T-cell therapy; 34 of 40 were HEp-2 negative and 6 of 40 were positive. **b**, Staining of all mAbs cloned from $EBV^-$ B cells obtained after CD19-CAR T-cell therapy; 9 of 10 were negative and 1 of 10 was positive. **c**, Controls: two mAbs cloned from healthy-control B cells (HC-5 and HC-9) and an anti-dsDNA positive control. Images were acquired under the conditions given for Supplementary Fig. 1. The 15% HEp-2 positivity of the pre-treatment $EBV^-$ repertoire is the background against which the $EBV^+$ repertoire in Fig. 3c should be read.

| Supplementary Table 1. Characteristics of 12 Patients Treated with CD19-CAR T Cells. | | | | | | | | | | | | |
|---|---|---|---|---|---|---|---|---|---|---|---|---|
| | **1** | **2** | **3** | **4** | **5** | **6** | **7** | **8** | **9** | **10** | **11** | **12** |
| **Demographic Characteristics** | | | | | | | | | | | | |
| Disease | SLE | SLE | SLE | SLE | SLE | SLE | SSc | SSc | SSc | IIM | IIM | IIM |
| Sex | female | female | male | female | male | Female | male | female | female | female | female | female |
| Age (years) | 24 | 43 | 21 | 25 | 29 | 55 | 56 | 29 | 23 | 81 | 45 | 56 |
| **Disease-specific Characteristics** | | | | | | | | | | | | |
| Disease Duration (years) | 5 | 1 | 1 | 2 | 5 | 25 | 4 | 4 | 1 | 2 | 1 | 1 |
| Disease Manifestations | Skin | Skin | Skin | Skin | Skin | Skin | Lung | Lung | Lung | Muscle | Lung | Lung |
| | Kidney | Joint | Kidney | Kidney | Kidney | Kidney | Skin | Skin | Skin | - | Muscle | Muscle |
| | Joints | Hema | Joints | Joints | Joints | Joints | - | - | - | - | - | - |
| | Serositis | - | - | Serositis | Serositis | - | - | - | - | - | - | - |
| | - | - | - | Hemato | - | - | - | - | - | -- | - | - |
| Autoantibody Entity | dsDNA | dsDNA | dsDNA | dsDNA | dsDNA | dsDNA | Scl70 | Scl70 | Scl70 | HMGCoA | MDA5 | Jo-1 |
| **Treatment Responses** | | | | | | | | | | | | |
| Duration of BC aplasia (days) | 168 | 86 | 55 | 203 | 153 | 181 | 84 | 57 | 84 | 84 | 170 | 84 |
| Naïve BC at reconstitution (%) | 96.2 | 96.5 | 98.8 | 98.7 | 97.7 | 98.4 | 97.4 | 97.9 | 95.1 | 97.7 | 97.9 | 97.8 |
| Memory BC at reconstitution (%) | 3.9 | 3.6 | 1.2 | 1.01 | 2.1 | 1.5 | 1.9 | 1.2 | 4.1 | 1.5 | 1.3 | 1.7 |
| Length of Drug-Free State (months) | 32.7 | 30.9 | 28.0 | 9.0* | 43.3 | 40.0 | 29.9* | 24.4 | 22.9* | 31.3 | 24.0 | 10.3* |
| Relapse | No | No | No | Yes | No | No | Yes | No | Yes | No | No | Yes |
| Reduction of autoantibody level | -100% | -99% | -98% | -100% | -100% | -87% | -79% | -70% | -53% | -100% | -100% | -72% |

| Supplementary Table 2. Characteristics of 8 SLE Patients Treated with Rituximab. | | | | | | | | |
|---|---|---|---|---|---|---|---|---|
| | **1** | **2** | **3** | **4** | **5** | **6** | **7** | **8** |
| **Demographic Characteristics** | | | | | | | | |
| Disease | SLE | SLE | SLE | SLE | SLE | SLE | SLE | SLE |
| Treatment | Rituximab | Rituximab | Rituximab | Rituximab | Rituximab | Rituximab | Rituximab | Rituximab |
| Sex | F | F | F | F | F | F | F | F |
| Age (years) | 61 | 31 | 42 | 25 | 62 | 25 | 30 | 23 |
| **Disease-specific Characteristics** | | | | | | | | |
| Disease Duration (years) prior to latest rituximab | >10 years | 11 | 21 | 6 | 17 | 3 | 0 | 9 |
| Autoantibodies | ANA, dsDNA, RNP, Ro | ANA, dsDNA, Sm, RNP | ANA | ANA, dsDNA, Sm, RNP, Ro | ANA, dsDNA, Ro | ANA, dsDNA, Sm, RNP | ANA, dsDNA, Sm, RNP | ANA, dsDNA |
| Prior Medications (before latest rituximab) | GC, HCQ, leflunomide, abatacept, belimumab, RTX | GC, HCQ, MTX, AZA, MMF, tacrolimus, RTX, CYC | GC, HCQ, AZA, MTX, MMF, belimumab, RTX | GC, HCQ, MMF, voclosporin, belimumab, CYC, RTX | GC, HCQ, MMF, AZA | GC, HCQ, CYC, AZA, mycophenolic acid, RTX | GC | GC, CYC, MMF, tacrolimus, RTX |
| **Treatment Responses** | | | | | | | | |
| Number of rituximab cycles pior to sampling | 2 | 2 | 2 | 2 | 1 | 3 | 1 | 2 |
| SLEDAI at time of post-Rituximab sample collection | 0 | 12 | 0 | 10 | 14 | 12 | 1 | 6 |
| Medications at time of post-rituximab sample collection | RTX | GC, MMF | HCQ, belimumab | GC, HCQ, MMF, voclosporin | GC | GC, MMF | GC | GC, HCQ, MMF, tacrolimus |

GC, Glucocorticoids; HCQ, Hydroxychloroquine; MTX, Methotrexate; AZA, Azathioprine; MMF, Mycophenolate mofetil; CYC, Cyclophosphamide; RTX, Rituximab

| **Supplementary Table 3. Individual patient characteristics and clinical response of the 7 patients with SLE treated with rituximab plus belimumab in the Synbiose-II trial.** | | | | | | | |
|---|---|---|---|---|---|---|---|
| **Patient** | 1 | 2 | 3 | 4 | 5 | 6 | 7 |
| **Study ID** | 11003 | 11005 | 11006 | 11023 | 11028 | 11059 | 11068 |
| **Demographic characteristics** | | | | | | | |
| **Age, years** | 41 | 42 | 17 | 46 | 40 | 21 | 38 |
| **Sex** | Male | Female | Female | Female | Female | Female | Female |
| **Race** | Other | White | White | White | White | Other | Black |
| **Disease duration, years** | 1 | 13 | 6 | 7 | 26 | 7 | 0 |
| **Organ involvement** | Nephritis, pericarditis | Nephritis | Nephritis, skin | Nephritis, arthritis, skin | Nephritis, arthritis, skin, leukopenia | Nephritis, NPSLE/lupus headache | Nephritis |
| **Lupus nephritis class** | II+V | IV | III | III+V | IV | III+V | V |
| **Prior therapies, N** | Pred, MMF | HCQ, MPS, pred, Aza, MMF | HCQ, MPS, pred, Aza, MMF | HCQ, Pred, MTX | HCQ, MPS, Pred, CYC, MMF, Aza, MTX, TAC | HCQ, Pred, MMF | HCQ, MPS, Pred, MMF, TAC |
| **Baseline autoantibodies** | | | | | | | |
| **Anti-dsDNA** | 1 | 1 | 1 | 1 | 1 | 1 | 0 |
| **Anti-Sm** | 0 | 0 | 1 | 0 | 0 | 0 | 0 |
| **Anti-RNP** | 0 | 0 | 1 | 0 | 0 | 0 | 1 |
| **Anti-SSA** | 0 | 1 | 0 | 1 | 0 | 1 | 0 |
| **Anti-SSB** | 0 | 0 | 0 | 0 | 0 | 1 | 0 |
| **Anti-C1q** | 0 | 0 | 0 | 1 | 0 | 1 | 0 |
| **Disease activity (SLEDAI)** | | | | | | | |
| SLEDAI, Baseline | 22 | 20 | 24 | 26 | 23 | 22 | 8 |
| SLEDAI, Wk 4 | 16 | 16 | 6 | 12 | 12 | 6 | 0 |
| SLEDAI, Wk 28 | 12 | 4 | 2 | 2 | 2 | 10 | 4 |
| SLEDAI, Wk 44 | 12 | 4 | 2 | 4 | 2 | 2 | 8 |
| SLEDAI, EoS | 8 | 4 | 2 | 4 | 2 | 2 | 2 |
| **Physician global assessment** | | | | | | | |
| PGA, Baseline | 3 | 1 | 3 | 2 | 3 | 3 | 3 |
| PGA, Wk 44 | 0 | 0 | 1 | 0.5 | 0 | 2 | 1.5 |
| PGA, EoS | 1 | 0 | 0.5 | 0 | 0 | 0 | 0 |
| **Concomitant therapy** | | | | | | | |
| Prednisone dose, Baseline | 7.6-20mg | 7.6-20mg | 7.6-20mg | 0mg | 0mg | 30-60mg | 7.6-20mg |
| Prednisone dose, Wk 4 | 20.1-30mg | 20.1-30mg | 20.1-30mg | 30-60mg | 7.6-20mg | 7.6-20mg | 20.1-30mg |
| Prednisone dose, Wk 28 | 0.1-5mg | 5.1-7.5mg | 5.1-7.5mg | 0.1-5mg | 0.1-5mg | 5.1-7.5mg | 5.1-7.5mg |
| Prednisone dose, Wk 44 | 0.1-5mg | 5.1-7.5mg | 5.1-7.5mg | 0mg | 0.1-5mg | 5.1-7.5mg | 5.1-7.5mg |
| Prednisone dose, Wk 104 | 0.1-5mg | 0.1-5mg | 0.1-5mg | 0mg | 0.1-5mg | 0.1-5mg | 0.1-5mg |
| Immunosuppression, Baseline | Pred 10, MMF | HCQ, pred , MMF | HCQ, pred 15, MMF | HCQ, MTX | MMF | HCQ, MMF | HCQ, Pred 40, MMF |
| Immunosuppression, Wk 4 | HCQ, BLM, MMF, Pred 40 | HCQ, BLM, MMF, Pred 40 | - | - | - | - | HCQ, BLM, MMF, Pred 40 |
| Immunosuppression, Wk 44 | HCQ,BLM, Pred 5mg | HCQ, BLM, Pred 7.5 | HCQ, BLM, Pred 7.5mg | HCQ, BLM | BLM, Pred 5mg | MMF, BLM, Pred 7.5mg | HCQ, BLM, Pred 7.5mg |
| **Clinical response** | | | | | | | |
| Clinical response, Wk 4 | yes | yes | yes | yes | yes | yes | yes |
| Clinical response, Wk 28 | yes | yes | yes | yes | yes | yes | yes |
| Clinical response, Wk 44 | yes | yes | yes | yes | yes | yes | no |
| Clinical response, EoS | yes | yes | yes | yes | yes | yes | yes |

| | | | | | | | |
|---|---|---|---|---|---|---|---|
| Clinical remission, Wk 4 | no | no | no | no | no | no | no |
| Clinical remission, Wk 28 | yes | no | no | yes | yes | no | no |
| Clinical remission, Wk 44 | yes | no | no | yes | yes | no | no |
| Clinical remission, EoS | yes | yes | yes | yes | yes | yes | yes |
| LLDAS, Wk 4 | no | no | no | no | no | no | no |
| LLDAS, Wk 28 | no | yes | yes | yes | yes | no | no |
| LLDAS, Wk 44 | no | yes | yes | yes | yes | no | no |
| LLDAS, EoS | no | yes | yes | yes | yes | yes | yes |
| Disease control, Wk 4 | no | no | no | no | no | no | no |
| Disease control, Wk 28 | no | no | yes | yes | yes | no | no |
| Disease control, Wk 44 | no | no | yes | no | yes | yes | no |
| Disease control, EoS | no | no | yes | no | yes | yes | yes |
| DORIS remission, Wk 4 | no | no | no | no | no | no | no |
| DORIS remission, Wk 28 | no | no | no | no | no | no | no |
| DORIS remission, Wk 44 | no | no | no | no | no | no | no |
| DORIS remission, EoS | no | no | no | no | no | no | no |
| **Treatment failure** | Non-adherence | - | - | - | Official non-response week 104 while response at week 100 due to no 50% decline in proteinuria, however ACEi stopped at week 100 | Insufficient reduction in proteinuria at week 28 (< 50% ) | Re-introduction of MMF and switch BEL to anifrolumab |
| **Week of treatment failure** | 28 | - | - | - | 104 | 28 | 44 |
| **B-cell reconstitution (CD19, cells/µl)** | | | | | | | |
| CD19, Baseline | 29 | 92 | 245 | 144 | 250 | 137 | 134 |
| CD19, Wk 4 | 43 | 355 | - | - | - | - | - |
| CD19, Wk 44 | 14 | 12 | 13 | 10 | 7 | 24 | 18 |
| **EBV-seq timepoints** | BL, W44 | BL, W44 | BL, W44 | BL, W44 | BL, W44 | BL, W44 | BL, W44 |

Response definitions are reported on all five criteria used in the trial. DORIS remission was not achieved by any patient in the belimumab + rituximab arm at any timepoint (0/7); LLDAS was reached by 6/7 at end of study.

All response states in this cohort are on-therapy states: patients remained on prednisone and/or maintenance immunosuppression throughout (see concomitant therapy rows). They are therefore not comparable to the drug-free remission reported for the CD19-CAR T cohorts.

EoS, end of study; LLDAS, Lupus Low Disease Activity State; DORIS, Definitions Of Remission In SLE; PGA, physician global assessment.
